# CheekAI: An explainable, deep learning epigenetic aging clock based on lifestyle and health epigenetic biomarker proxies

**DOI:** 10.64898/2026.11.21.768329

**Authors:** Maxim N. Shokhirev, Michael J. Corley, Varun B. Dwaraka, Adiv A. Johnson

## Abstract

The human methylome is poorly understood and, as such, the vast majority of epigenetic aging clocks are unexplainable black boxes. To address this, we created CheekAI, a robust deep learning model that uses lifestyle and health epigenetic biomarker proxies (EBPs) to predict age. Trained using a longitudinal dataset of 25,400 adult buccal DNA methylation samples derived from 14,716 distinct adults, we used principal components and multi-objective optimization to produce a clock that is explainable, reproducible, responsive, and accurate. The Pearson correlation with chronological age is 0.969, the mean absolute error is 2.49 years, and the intraclass correlation coefficient across 679 replicate pairs is 0.990. CheekAI was significantly associated with a plethora of lifestyle and health variables, including resting heart rate, omega-3 intake, ultra-processed food intake, daily step count, resistance training, and time spent in nature. When applied to external saliva and buccal datasets, CheekAI significantly captured a variety of signals, including smoking and oesophageal cancer. CheekAI was also significantly reduced by a multifactorial lifestyle intervention and this reduction was driven by improvements in dietary and metabolic EBPs. This novel artificial intelligence model represents the first instance of a buccal epigenetic aging clock that utilizes EBPs to offer interpretability and actionability.

## Introduction

The human methylome [1] encompasses around 28 million DNA methylation sites in the approximately three billion base pair genome. This layer of data is, on the whole, quite enigmatic. While methylomic information can be used to predict a wide array of variables [2, 3], how changes in methylation lead to alterations in protein expression or functional changes is poorly understood. This presents a fundamental challenge for epigenetic aging clocks, a newer class of computational biomarkers that predict age using DNA methylation data as inputs [4].

To date, the vast majority of epigenetic aging clocks are difficult-to-interpret black boxes [5]. This means that, in an interventional setting, it is challenging to explain why a given clock was shifted upwards or down. Similarly, black box models cannot offer coherent explainability or actionability. This is quite relevant to highlight given that both explainability and actionability have been documented to be highly desirable qualities for an aging biomarker [6]. If a cohort is predicted to have an older epigenetic age, a non-explainable model is not able to offer guidance on why epigenetic age is elevated or what to do about it. Similarly, traditional models cannot satisfactorily explain why disparate clocks provide distinct results [7].

Recently, explainable artificial intelligence (XAI) models have emerged that inject interpretability into aging clocks [8, 9]. Epigenetic XAI models do this by either abstracting DNA methylation sites to epigenetic biomarker proxies (EBPs) – methylation-based predictors of non-age variables like proteins, metabolites, clinical variables, and fitness metrics – or to pathways or networks via annotated genes. The deep learning model EpInflammAge, for instance, uses methylation-based predictors of circulating cytokines and chemokines as inputs for age prediction [10]. Conversely, the models EXP.REACTOME [11] and XAI-AGE [12] abstract CpG sites linked to annotated genes to Reactome [13] pathways and utilize these to estimate age.

Inspired by the advent of principal component (PC)-based clocks [14] as well as the specific next-generation models CheekAge [15], OMICmAge [2], and SystemsAge [16], here we develop a new XAI buccal clock called CheekAI. CheekAI is a highly reproducible deep learning model that uses PCs and EBPs to make interpretable, reproducible age predictions in adult buccal data. To date, this represents the first buccal clock that utilizes EBPs.

## Results

### Cohort and model design

The previously described CheekAge model was trained on a cohort of 8,045 adults [15]. Years later, that private adult buccal database has dramatically expanded and become longitudinal in nature. Here, we analyze 25,400 adult buccal DNA methylation profiles (53.5% male, 46.5% female) from 14,716 distinct adults (Figure 1A) in this database that were collected across three Illumina methylation microarray platforms (Figure 1B). Specifically, a total of 11,852, 9,609, and 3,939 samples correspond to an EPICv1, an EPICv2, or a custom EPIC-based array, respectively. The latter custom array contains close to 250,000 DNA methylation sites. A total of 679 replicates contribute to this dataset (Figure 1B), 556 of which are across platforms and 123 of which are within the same platform. 3,402 people have at least two timepoints and more than 2,000 have at least three timepoints spanning a period of time greater than six months (Figure 1C). Every buccal sample was paired with up to 50 self-reported or measured lifestyle, health, demographic, and technical variables (Figure 1D). Much of this was collected via a digital survey that asked questions about diet, fitness, sleep, mental health, and habits.

**Figure 1.**
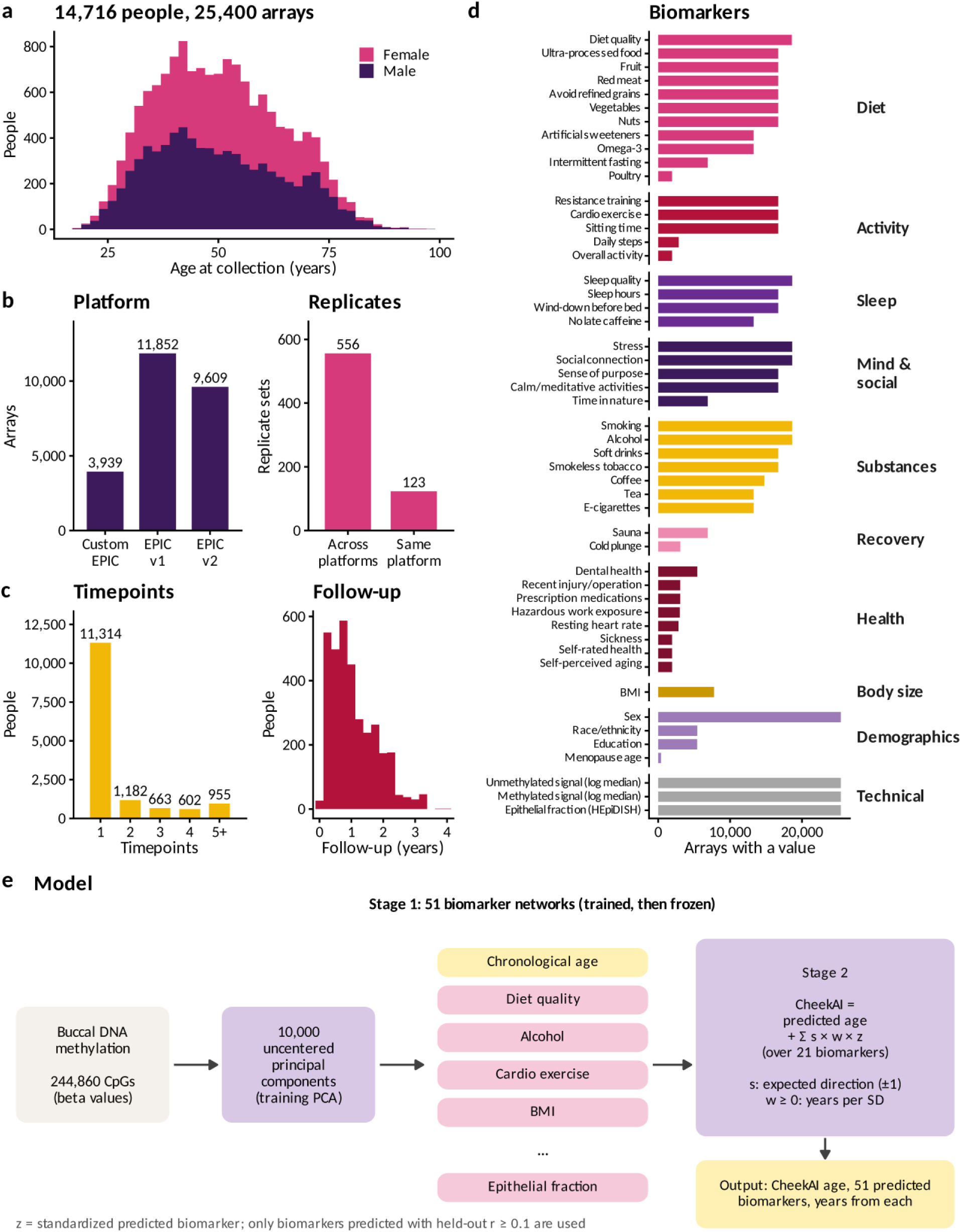
Cohort, biomarkers and model. (a) Age at collection by sex, counting one sample per person. (b) Number of arrays measured on each platform, and the replicate sets whose arrays were measured on one platform or on more than one. (c) Number of timepoints per person, and the follow-up time of people sampled more than once. (d) The 50 biomarkers modeled besides chronological age, grouped by domain, with the number of arrays that have an observed value for each. (e) The CheekAI model. Beta values were projected onto 10,000 principal components of the training data. In stage 1, 51 networks are trained separately and then frozen; they predict chronological age and 50 lifestyle, health, demographic, and technical biomarkers. In stage 2, CheekAI is the predicted chronological age plus, for each of the 21 biomarkers that pass the validity filter, a sign-constrained effect in years per standard deviation of the standardized predicted biomarker.

At a high level, CheekAI was trained via multi-objective optimization to produce a clock that offers explainability, reproducibility, accuracy, and responsiveness. Analogously to many next-generation clocks that are trained via a multi-stage approach, CheekAI was built in two stages (Figure 1E). Given that PCs are known to suppress technical noise [14], we first reduced methylation profiles to 10,000 PCs, from which 51 separate neural networks predict either chronological age or 50 EBPs (made possible via the aforementioned metadata). These networks were trained and then frozen. From here, CheekAI represents the predicted chronological age plus a sum of 21 per-biomarker adjustments, each of which is constrained to the direction in which the biomarker coherently relates to aging. For instance, a healthier predicted diet can only lower CheekAI. The size of each adjustment is learned by optimizing four objectives jointly: accuracy, agreement between replicates, age-adjusted association with the observed lifestyle factors, and agreement between within-person changes in epigenetic age and lifestyle over time. Because the adjustment is additive, every CheekAI value decomposes exactly into predicted age plus a contribution, in years, from each biomarker. All evaluations use person-grouped nested cross-validation, meaning that no person contributes to both training and evaluation and every modeling choice is made within the training cohort.

### Accuracy and reproducibility

On held-out individuals, CheekAI was highly correlated with chronological age (Figure 2A) with a Pearson correlation of 0.969, a mean absolute error (MAE) of 2.49 years, and a root mean square error (RMSE) of 3.46 years. Across 679 replicate pairs, CheekAI was highly reproducible with an intraclass correlation coefficient (ICC) of 0.990, a mean absolute replicate difference of 1.41 years, and a median absolute replicate difference of 1.11 years (Figure 2B). This is on par with what has been previously reported for PC clocks in buccal replicates [17].

**Figure 2.**
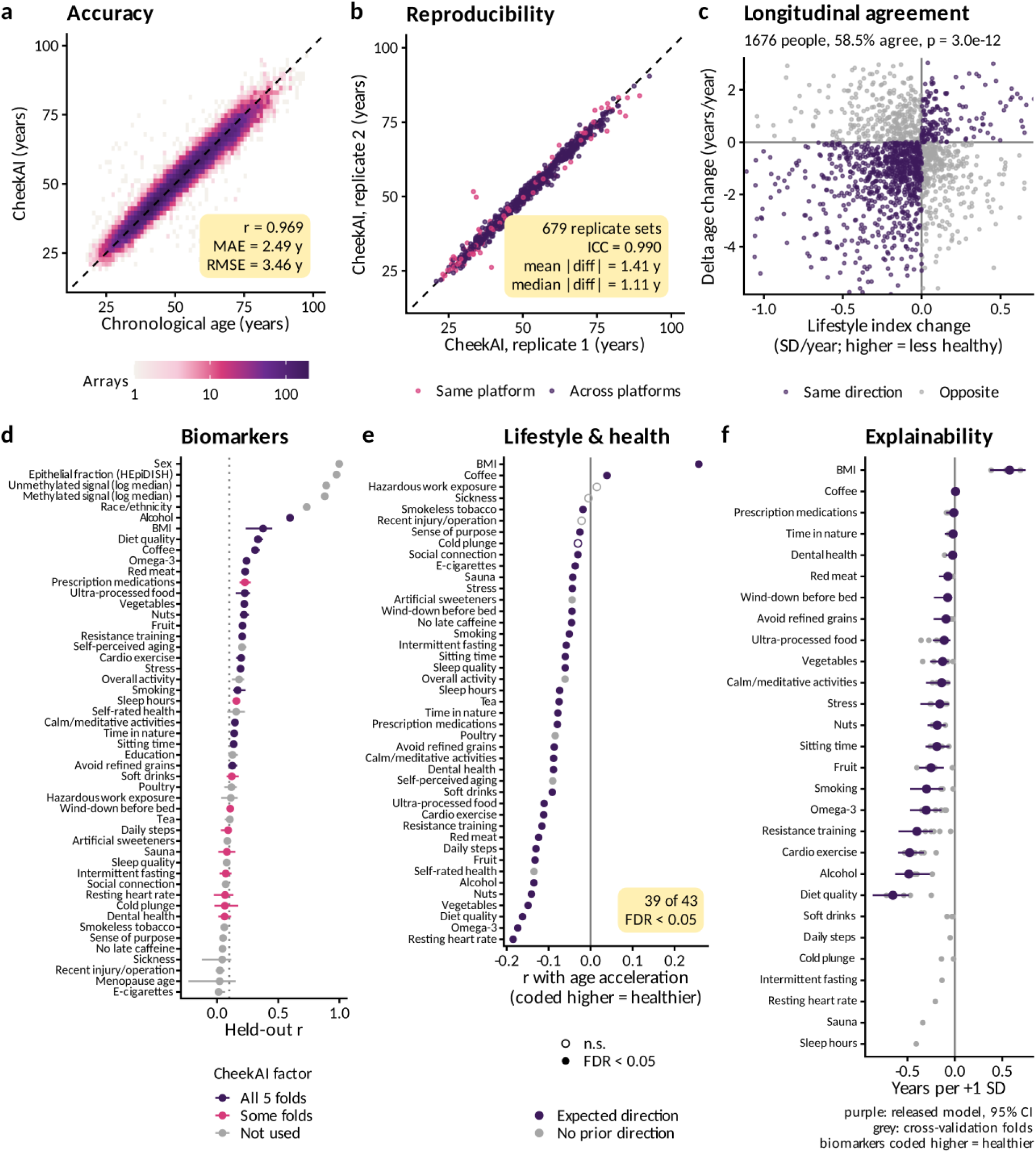
Performance of CheekAI on held-out individuals. All panels pool the five outer folds of person-grouped cross-validation, so every array is predicted once by a model that never saw that person. (a) CheekAI against chronological age. (b) CheekAI of the first two arrays of each technical-replicate set (679 sets). Pink points are pairs measured on the same platform and purple points pairs measured on different platforms. (c) For each person with at least three timepoints spanning more than six months, the change per year in platform-corrected CheekAI delta age (epigenetic age – chronological age) against the change per year in the standardized, sign-reversed observed lifestyle index, where higher values are coded as less healthy. People whose two changes share a sign are shown in purple and the rest in grey, and the axes show the central 96% of people. (d) Held-out correlation of each stage-1 predicted biomarker with its observed value. Points show the mean over folds and lines the range across folds. The dotted line marks the validity threshold of r = 0.1. Purple biomarkers entered stage 2 in all five folds, pink biomarkers in some folds, and grey biomarkers in none (this includes the demographic and technical variables). (e) Correlation of CheekAI delta age with each observed lifestyle and health factor. Filled points are significant at FDR < 0.05 and open points are not. Purple factors have an expected direction from training people, and every significant association with such a factor was in that direction; grey factors have no expected direction. Survey answers are coded so that higher values are presumably healthier. (f) Years added to CheekAI per +1 standard deviation of each predicted biomarker in the released model, with bootstrap 95% confidence intervals in purple. Grey dots show the same effect estimated in each cross-validation fold. The factors at the bottom were used in some folds but not in the released model, so they have fold estimates only.

### Longitudinal agreement

One piece of metadata that is unique to this dataset is a lifestyle index (with a maximum score of 100 points) based on self-reported lifestyle behaviors collected via a digital survey.

People with healthier behaviors (e.g., those who reported engaging in more frequent exercise) received higher scores while those who engaged in less healthy behaviors (e.g., those that reported smoking or frequently consuming ultra-processed food) garnered lower scores. A total of 1,676 people had a lifestyle score at three or more timepoints spanning a period greater than six months. The direction of change in CheekAI delta age (which is epigenetic age – chronological age) aligned with the direction of change in the lifestyle score in 58.9% of individuals (binomial p = 3.5 x 10^-12^) and in 58.5% after removing platform offsets estimated in the training group (p = 3.0 x 10^-12^). Put differently, individuals who reported improvements to their lifestyle over time were more likely to have a younger CheekAI when tested again in the future (Figure 2C).

### Explainable biomarkers

Stage-1 networks (Figure 1E) predicted 50 non-age biomarkers (spanning lifestyle, health, demographic, and technical variables) with a median held-out correlation of 0.14 (Figure 2D). Using a pre-specified validity filter of a held-out r ≥ 0.1 and a false discovery rate (FDR) < 0.05, 18 lifestyle and health biomarkers passed in every fold and 28 passed in at least one fold. The best EBPs were alcohol intake (r = 0.60), body mass index (BMI) (0.38), diet quality (0.33), and coffee intake (0.31). The technical and cellular variables were also accurately predicted (e.g., epithelial fraction r = 0.98).

CheekAI delta age was associated with 39 of the 43 lifestyle and health factors at an FDR < 0.05, with a mean absolute Pearson correlation of 0.084 (Figure 2E). For the vast majority of these, the directionality was intuitive and what one would expect. For instance, consuming more vegetables, engaging in more cardiovascular exercise, performing more strength training, minimizing ultra-processed foods, spending more time in nature, and not smoking were all associated with a younger epigenetic age (Figure 2E).

The released CheekAI model, which is trained on all samples, uses a total of 21 biomarkers (Figure 2F), which are: BMI, daily coffee intake, number of prescription medications, time spent in nature, self-assessed dental health, red meat intake, whether or not someone winds down before bed, intake of refined grains, ultra-processed food intake, vegetable intake, engagement in calm or meditative activities, perceived stress, nut intake, time spent sitting each day, fruit intake, smoking status, omega-3 fatty acid intake (from any source), weekly resistance training, weekly cardiovascular exercise, weekly alcohol intake, and overall diet quality. As an example of how this works, CheekAI changes by -0.65 years for diet quality with a bootstrap 95% confidence interval (CI) of -0.87 to -0.49 per one standard deviation (SD) shift of the predicted biomarker. CheekAI similarly changes by −0.48 for alcohol intake (−0.63 to −0.26), −0.48 for cardiovascular exercise (−0.60 to −0.33), −0.16 for stress (−0.36 to −0.06), and +0.58 for BMI (0.37 to 0.75). The stability of CheekAI across cross-validation folds is visualized in Supplementary Figure S1.

### Comparison with CheekAge

Using the same held-out samples, we compared CheekAI to the previously described next-generation epigenetic aging clock CheekAge (Figure 3). Unlike CheekAI, CheekAge is a non-explainable, non-deep learning model trained on cross-sectional data [15]. In terms of accuracy with chronological age (Figure 3A), CheekAge had a Pearson correlation of 0.958 (compared to 0.969 for CheekAI), an MAE of 3.23 years (compared to 2.49 years for CheekAI), and an RMSE of 4.25 years (compared to 3.46 years for CheekAI). A non-explainable stage-1 age network (Figure 1E) was slightly more accurate (r = 0.982, MAE = 1.70 years, and RMSE = 2.65 years), indicating that optimizing for other parameters results in a slight accuracy tradeoff for both CheekAge and CheekAI (Figure 3A). In terms of reproducibility (Figure 3B), CheekAge was also less reproducible than CheekAI with a replicate ICC of 0.982 (compared to 0.990 for CheekAI). Longitudinal agreement was also relatively lower for CheekAge (Figure 3C). Specifically, CheekAge showed longitudinal agreement of 55.0% (p = 4.5 x 10^-5^) that dropped to 50.9% (p = 0.48) after platform correction. While CheekAI was significantly associated with 39 different survey factors with a mean absolute Pearson correlation of 0.084, CheekAge was associated with 37 distinct variables with a mean absolute Pearson correlation of 0.058 (Figure 3D). Thus, CheekAI was more accurate, more reproducible, more responsive, and more strongly associated with holistic lifestyle and health factors than CheekAge in this dataset.

**Figure 3.**
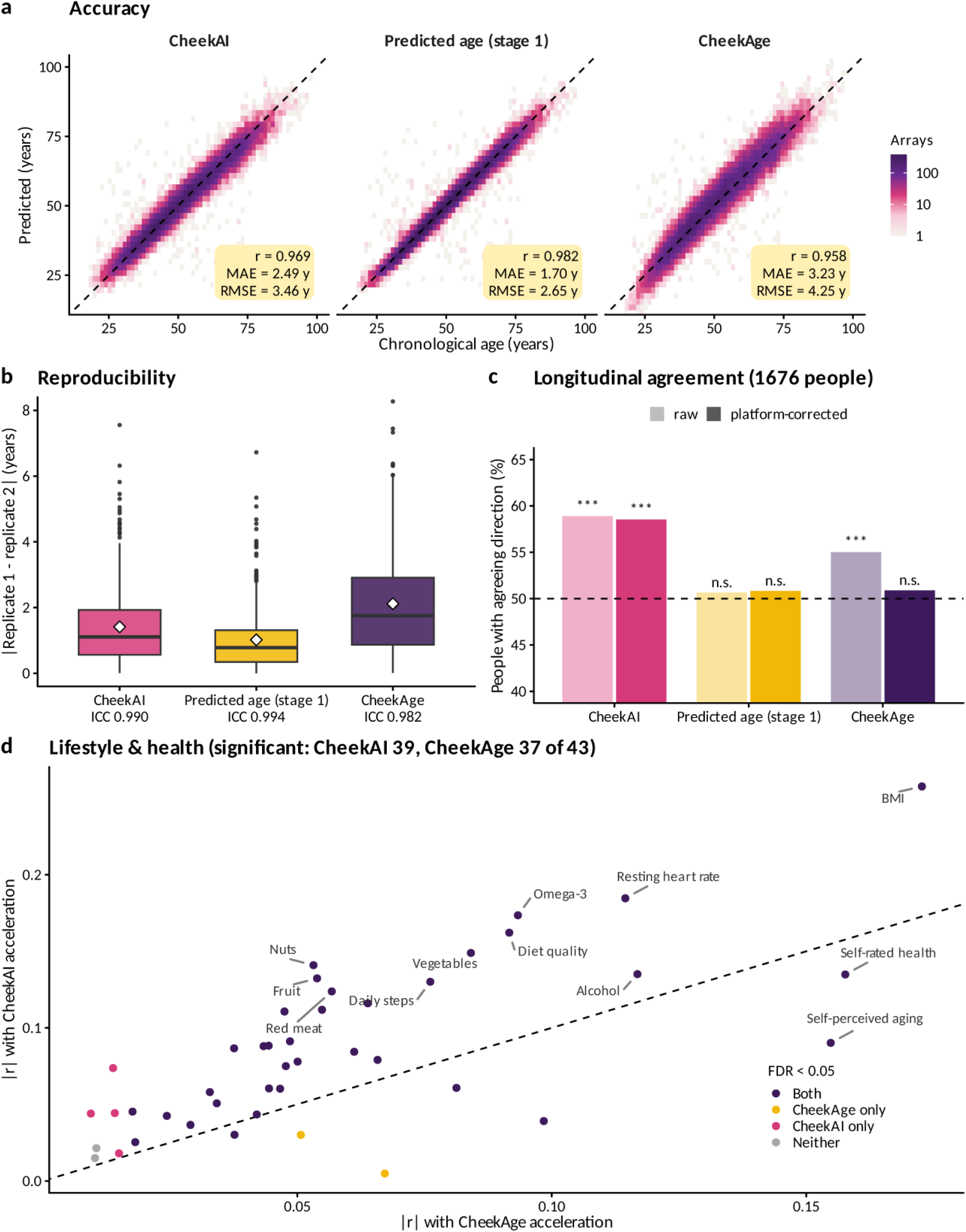
Comparison with CheekAge. CheekAI, predicted chronological age from stage 1 alone, and CheekAge on the same held-out arrays. (a) Accuracy against chronological age for each clock. (b) Absolute difference between the first two arrays of each technical-replicate set; boxes show the quartiles, diamonds the mean, and the one-way ICC is given under each clock. (c) Longitudinal agreement before (light bars) and after (dark bars) removing platform offsets estimated in training people. The dashed line marks the 50% expected by chance; three asterisks denote binomial p < 0.001 and n.s. denotes not significant. (d) Absolute correlation of each factor with CheekAI age acceleration against its absolute correlation with CheekAge age acceleration, colored by which clock, if either, is significantly associated with the factor at FDR < 0.05. The dashed line is the identity, so points above it are factors more strongly associated with CheekAI.

CheekAge was trained on an earlier subset of this cohort that contained around 8,000 samples, most of which are part of the held-out arrays. Since these arrays were held out from CheekAI but may have been seen by CheekAge, we restricted the analysis to the arrays from people that neither clock was trained on. When doing so, the comparative results were similar (Supplementary Table S1). CheekAI had an RMSE of 3.23 years vs. 4.24 years for CheekAge, a replicate ICC of 0.992 vs. 0.985 across 346 replicate sets (a mean difference of 1.27 years vs. 1.99 years), and 34 vs. 29 of 38 testable associations with a mean absolute Pearson correlation of 0.086 vs. 0.049. Its longitudinal agreement was 58.7% vs. 52.2% after platform correction (p = 3.0 x 10^-10^ vs. 0.12). Looser reproducibility limits (ICC ≥ 0.985, replicate error ≤ 1.5 y) gave stronger associations (mean |r| = 0.098) at the cost of accuracy (RMSE = 4.07 years) and reproducibility (ICC = 0.986) with similar longitudinal agreement of 58.9% (Supplementary Table S2). Sample size estimates for CheekAI and CheekAge across various interventional trial designs and effect sizes are summarized in Supplementary Table S3. These can be interpreted as the number of samples needed to detect a significant signal relative to noise in an interventional setting.

### Validation in independent, public datasets

Similarly to before [18], we explored the performance of CheekAI across publicly available methylomic datasets. Specifically, we analyzed 12 DNA methylation datasets in the Gene Expression Omnibus [19] and ArrayExpress [20] databases that compared people who differed in lifestyle exposure, intervention, or disease status (see Methods). Because CheekAI is an XAI model dependent on EBPs, the contribution of each EBP to an association can be explored.

In the 450K buccal dataset GSE94876 containing 40 cigarette smokers and 40 non-tobacco users [21], delta CheekAI was 1.72 years higher in smokers with a p-value of 0.04 (Figure 4A). The biomarkers contributed 0.91 years (p = 0.005) and predicted age contributed 0.81 years (p = 0.21). The smoking EBP was the largest of the 21 terms (+0.21 years, FDR = 0.028) followed by alcohol (+0.12 years, FDR = 0.028). The predicted smoking biomarker separated smokers from non-smokers with an area under the curve (AUC) of 0.69. This held even though 45% of the CpGs used by CheekAI are not on the 450K array.

**Figure 4.**
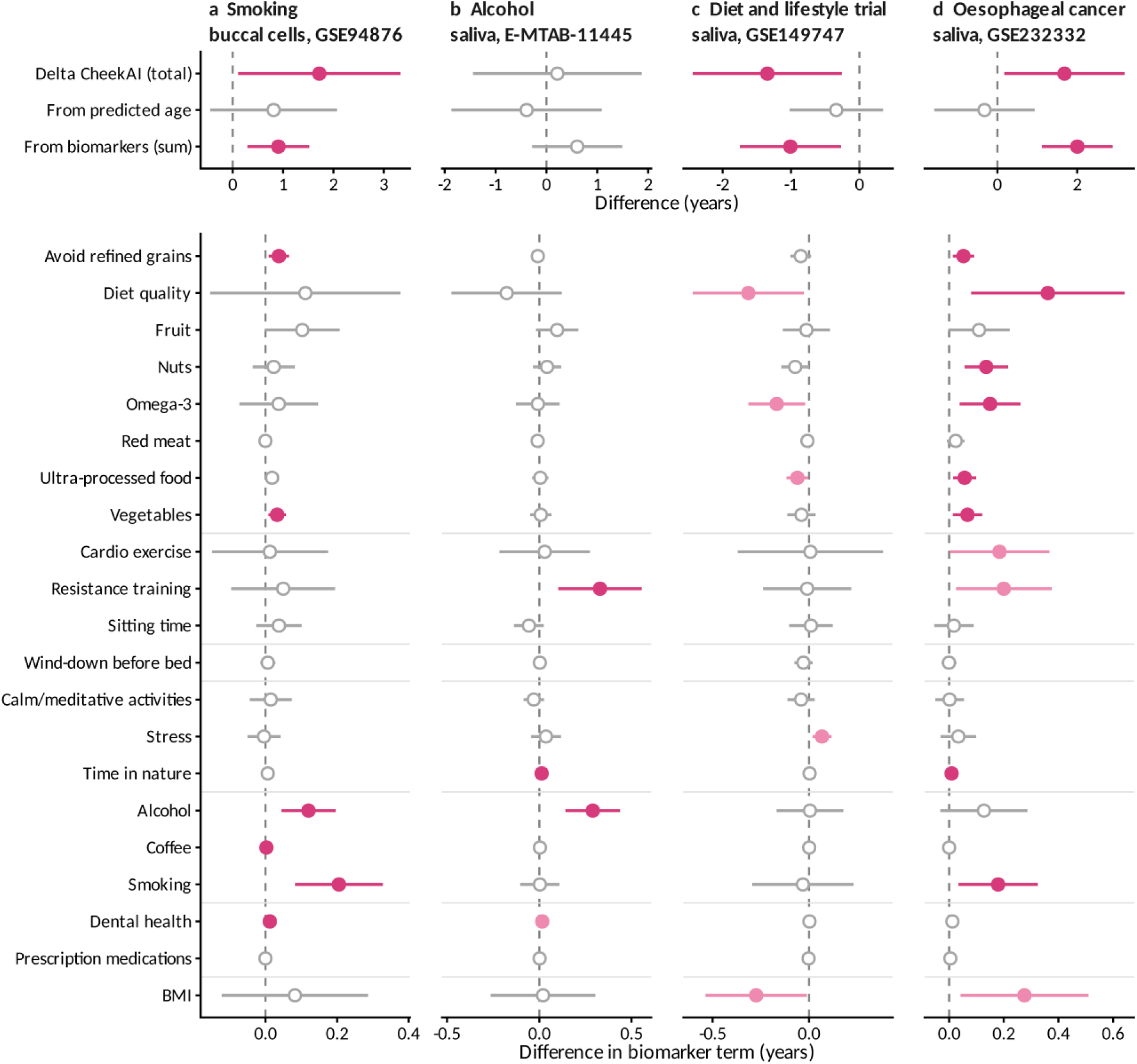
CheekAI in independent public datasets. Each column is one dataset, analyzed with CheekAI. (a) Buccal cells from cigarette smokers and non-tobacco users (GSE94876). (b) Saliva from Swedish adults who reported drinking alcohol on many days or a few days each week (E-MTAB-11445). (c) Saliva from a randomized trial of an eight-week diet and lifestyle program involving treatment and control groups (GSE149747). Each value is the change over the eight weeks in the treatment group minus the change in the controls, so negative values mean that CheekAI fell more in the treated individuals. (d) Saliva from people with oesophageal adenocarcinoma or high-grade dysplasia and controls (GSE232332). For each dataset, the top rows show the group difference in delta CheekAI (CheekAI adjusted for age within the dataset) and its exact split into the part from predicted age and the part from the biomarkers, which is the sum of the 21 biomarker terms. The bottom rows show the same difference for each biomarker term, grouped by domain and in the same order for every dataset. Positive values mean older in the exposed or case group. Points are estimates, lines are 95% confidence intervals, and each column has its own x axis. Filled dark pink points are significant, at p < 0.05 for the top rows and at Benjamini– Hochberg FDR < 0.05 across the 21 terms for the bottom rows. Filled light pink terms are significant at p < 0.05 before but not after this correction, and open grey points are not significant. Some significant terms lie close to zero because their biomarker adds little to CheekAI.

In a larger saliva EPIC dataset E-MTAB-11445 derived from Swedish adults [22], the alcohol EBP was 0.29 years higher in people who reported drinking several days each week than in those who reported drinking less (Figure 4B). The FDR value for this difference was 0.003 and alcohol was the largest of the 21 EBPs. The predicted alcohol biomarker also separated the two groups with an AUC of 0.65 and, conversely, the smoking term did not significantly differ (p = 0.96). Resistance training (+0.33 years) and time in nature (+0.01 years) also differed at an FDR < 0.05. Interestingly, the alcohol-based difference was too small to move the entire delta CheekAI score as a whole (+0.22 years, p = 0.80).

Delta CheekAI in saliva was significantly reduced by a randomized trial involving an eight-week, multifactorial lifestyle-based program in dataset GSE149747 (Figure 4C) [23]. This trial involved both interventional (n = 18) and control (n = 20) groups. Delta CheekAI was reduced by 1.34 years in treated individuals compared to controls (p = 0.021). This was driven by the diet quality (-0.32 years), omega-3 intake (-0.17 years), ultra-processed food intake (-0.06 years), and BMI (-0.27 years) EBPs. Interestingly, the stress EBP moved up (+0.07 years). In total, all 21 EBPs carried 1.01 years of the overall reduction in CheekAI (p = 0.011). In the separate saliva dataset GSE232332 involving 98 people with oesophageal adenocarcinoma or high-grade dysplasia and more than 100 controls [24], delta CheekAI was 1.68 years higher in cases (p = 0.03). This elevation was driven by changes to eight EBPs (Figure 4D), including smoking (+0.18 years, FDR = 0.043) and diet quality (+0.36 years, FDR = 0.043).

The additional datasets analyzed are summarized in Supplementary Table S4. These results, which are mixed, span a variety of tissues and health variables. Of note, delta CheekAI was 0.89 years higher in blood (p = 0.007) derived from patients with current post-traumatic stress disorder in the dataset GSE132203 [25]. This elevation, however, came from predicted age and not the EBPs.

### Assessing the impact of semaglutide

Recently, a post hoc analysis of the effects of the GLP-1 receptor agonist semaglutide on epigenetic biomarkers in patients with HIV-associated lipohypertrophy was published [26]. The actual dataset came from a trial involving treatment (n = 45) and placebo (n = 39) groups. Looking at the blood methylomic datasets from those trial participants, we see that the predicted age component of CheekAI was reduced (Supplementary Figure S2A). While the overall CheekAI model was decreased, this drop did not reach significance. The majority of the EBPs were statistically unchanged, which indicates that buccal EBPs cannot be cleanly translated to blood samples. All other tested epigenetic aging clocks that were trained using buccal samples were not significant (Supplementary Figure S2B). These findings suggest that CheekAI and other buccal clocks are more appropriate for cheek swab or saliva datasets and may not work as well in other tissue types.

## Discussion

Inspired by a variety of previously published models, here we create the first-ever XAI buccal clock based on EBPs. To our knowledge, this is the first paper to demonstrate that such a breadth of lifestyle-oriented EBPs can be constructed based on digitally collected survey data. Unlike CheekAge, which utilizes more than 200,000 CpGs and is a black box, CheekAI can be explained by the status of the dozens of EBPs that feed into a final age prediction.

One of the first things we did was explore how the CheekAI epigenetic age score was associated with 43 different lifestyle and health variables. Intriguingly, one of the metrics with the strongest correlations with CheekAI was omega-3 fatty acid intake. This variable came from a question that asked people how many days a week they consume omega-3 fatty acids from any source. The fact that this emerged as one of the most robust associations is fascinating given that various next-generation epigenetic aging clocks have been reduced by omega-3 fatty acid supplementation [27]. One especially robust trial involving 777 healthy adults showed that omega-3 supplementation for more than three years significantly lowered DunedinPACE, GrimAge2, and DNAm PhenoAge [28]. More broadly, multiple studies have identified a correlation between epigenetic aging clocks and omega-3 intake [29-31]. The omega-3 fatty acid alpha-linolenic acid has also been reported to extend lifespan in worms [32]. More research is needed to better understand the relationship between omega-3 fatty acids, DNA methylation, and aging.

In independent saliva and buccal validation datasets, it was notable to see that the EBP changes were quite coherent. In buccal cells derived from smokers, the most dramatically elevated EBP was for smoking. Likewise, the alcohol EBP was significantly elevated in saliva samples derived from individuals that consumed more alcohol. In the trial involving a multifactorial lifestyle intervention (which included dietary changes), the significant drop in CheekAI was mediated by improvements in the diet quality, omega-3 fatty acid intake, ultra-processed food intake, and BMI EBPs. While none of the EBP changes were significant on their own, together they helped create a significant drop in the overall epigenetic age estimate. The stress EBP was also mildly elevated in response to this putatively beneficial intervention. This is worth noting because many beneficial adaptations appear to induce a period of epigenetic stress and remodeling. This has, for example, been observed for intermittent hypoxia in mice [33] and sauna use in humans [34].

Taken together, our work demonstrates that it is possible to create an XAI buccal clock that accurately correlates with chronological age, offers reproducibility across replicates, shows an ability to respond in an interventional setting, and has the capacity to associate with clinically relevant variables. More broadly, this multi-optimization deep learning approach may be a promising strategy for creating future next-generation models. We also show, for the first time, that EBPs can be created for a wide variety of digitally collected lifestyle variables. Additional work is warranted to see if more robust explainability can be offered, such as by generating EBPs with higher Pearson correlations with the metrics they estimate.

## Methods

### Cohort and data

Buccal swabs from 14,716 adults (25,400 arrays) were profiled on Illumina EPIC v1, EPIC v2 or a custom EPIC-based array. IDAT files were processed with minfi [35] using noob background and dye-bias correction [36]. Subsequent analyses used the 244,860 CpGs measured on all three platforms. Each sample had chronological age and up to 50 lifestyle, health, demographic, and technical variables. Survey answers relevant to diet, activity, sleep, mind C social, substances, recovery, health, body size, demographics, or technical components were coded so that higher values are healthier. The epithelial cell fraction was estimated with EpiDISH [37], and log median methylated and unmethylated signal intensities from minfi. Technical replicates and repeat collections from the same person were linked from the sample metadata and people were defined as the connected groups of linked arrays.

### Stage 1: biomarker networks

Beta values were represented by 10,000 uncentered PCs, computed exactly from the sample Gram matrix and refitted inside every training set so that held-out samples never informed the components. New samples are projected onto the stored training loadings. For each of the 51 targets (chronological age and 50 biomarkers), a separate feed-forward network was trained on the PCs in PyTorch [38] with the Adam optimizer [39]. The loss was a class-weighted mean squared error plus a penalty on disagreement between technical replicates. The learning rate, weight decay and stopping epoch were chosen per biomarker by grid search on an inner validation fold. The networks were then frozen.

### Stage 2: CheekAI

CheekAI is defined as:

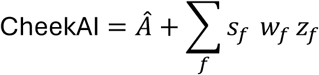

where *Ã* is the predicted chronological age, *z*_*f*_is the standardized predicted value of biomarker *f*, *w*_*f*_ ≥ 0 is a learned effect in years per SD, and *s*_*f*_ ∈ {−1, +1} is the expected direction of the biomarker. The expected direction is the sign of the association between the observed factor and CheekAge [15] acceleration in the training people, so a healthier predicted value can only lower CheekAI. Only lifestyle and health biomarkers predicted with an out-of-fold correlation of at least 0.1 (Benjamini–Hochberg FDR < 0.05) entered stage 2. The weights minimize a weighted sum of four terms: the error against chronological age, the squared disagreement between replicate arrays, the shortfall from perfect direction-oriented correlation between age acceleration and the observed factors, and the shortfall from perfect correlation between per-person slopes over time of age acceleration and of a survey-based lifestyle index. The four loss weights were chosen from a grid of 60 settings by inner cross-validation with a rule fixed before evaluation: among settings with a one-way replicate ICC of at least 0.99 and a mean replicate difference of at most 1.25 years, the setting with the largest sum of the association and longitudinal terms was selected.

### Validation

All reported results come from person-grouped nested cross-validation with five outer and four inner folds, so no person contributed to both training and evaluation. All modeling choices, including the principal component analysis (PCA), the stage-1 settings, the validity filter, the expected directions and the stage-2 weights, were made within the training people of each outer fold. Each array was then predicted once by a model that had never seen that person. The released model was trained on all 25,400 arrays with the same procedure.

### Evaluation

Accuracy was measured by the correlation, MAE, RMSE and mean bias against chronological age. Reproducibility was measured by the one-way ICC [40], and the absolute difference between the first two arrays of each technical-replicate set. The ICC was also computed on the delta age scale with a linear mixed model in lme4 [41]. Age acceleration was defined as the residual of the difference between a clock and chronological age, regressed on chronological age within each held-out fold. Its Pearson correlation with each observed factor was tested with Benjamini–Hochberg control of the FDR [42]. For longitudinal agreement, people with at least three timepoints spanning more than six months were scored on whether the slope of their delta age and the slope of their observed lifestyle index had the same sign, with a two-sided binomial test against 50%. This was done with and without subtracting platform offsets estimated in training people. Confidence intervals for the effects came from 100 person-level bootstrap refits of stage 2. CheekAge was computed with the published model for each array’s platform and evaluated with the same code on the same held-out arrays. Because CheekAge was trained on about 8,000 arrays from this cohort, all comparisons were repeated on the arrays with a “TH” barcode, none of which was used to train CheekAge.

### Publicly available external datasets

We searched Gene Expression Omnibus [19] and ArrayExpress [20] for DNA methylation datasets from buccal cells, saliva, or blood measured on the Illumina 450K or EPIC array, with at least 50 samples and a lifestyle exposure, intervention, or disease. IDAT files were processed with minfi noob when they were available. Otherwise the deposited beta values were used. CheekAI was predicted and CpGs missing from a dataset were set to their training means. Delta CheekAI was the residual of CheekAI on chronological age within the dataset, and it splits exactly into the residual of predicted age and the biomarker part, the sum of the 21 biomarker terms. Group differences in each of these and in each biomarker term were estimated by linear regression, with age and sex as covariates where they varied. Paired designs used the change within each person between timepoints, with repeated arrays averaged. Because the models are linear, the effect on delta CheekAI is exactly the sum of the effects on its two parts. P values for the 21 terms were corrected with the Benjamini–Hochberg method within each test.

The Extended Methods provide additional methodological information, including details relevant to sample metadata and biomarker coding, the PCA computation and out-of-sample projection, the network architectures and hyperparameter grids, the handling of collapsed fits and numerical precision, the stage-2 loss and weight grid, the cross-validation design, the metric definitions, the CheekAge comparison, the analysis of each public dataset, and the software used.

### Semaglutide randomized trial analysis

We evaluated CheekAI in paired baseline and week-32 peripheral blood samples from a previously published randomized, placebo-controlled semaglutide trial [26]. Corrected participant and visit annotations identified 84 complete pairs, comprising 45 participants assigned to semaglutide and 39 assigned to placebo; non-randomized controls were excluded. Frozen predictors were applied to source-normalized, ssnoob-processed EPICv2custom methylation data without retraining, recalibration, or refitting the methylation PCs. CheekAI inputs included 232,895 of 244,860 required CpGs (95.1%); absent sites used the model’s fixed external imputation values.

For CheekAI, its predicted-age component, the sum of its biomarker contributions, and each of the 21 weighted biomarker terms, we regressed week-32 minus baseline values on randomized arm, baseline composite CheekAI, baseline age, and sex. The same participants and design matrix were used for all outcomes, preserving additivity of the estimated treatment effects. For the clock comparison, we evaluated CheekAI, its age-only network, CheekAge [15], Horvath’s multi-tissue clock [43], the Skin C Blood clock [44], and the PC versions of the Horvath multi-tissue and Skin C Blood models [14]. Each clock model adjusted for its own baseline score, baseline age, and sex. Consequently, the age-network coefficient in the clock comparison differs slightly from the corresponding component coefficient in the additive analysis.

Treatment coefficients represent the adjusted difference in change for semaglutide relative to placebo, in years. We used HC3 standard errors, two-sided normal-approximation p values, and nominal 95% confidence intervals. Benjamini–Hochberg correction was applied separately across the biomarker terms and clock outputs. Week-20 results and cell-composition, recorded-shipment, elapsed-time, and chip sensitivities were retained in the source tables, including non-estimable fits. Inference concerns participants with complete pairs.

## Data Availability

The following publicly listed datasets were utilized in this study: GSE94876, GSE149747, GSE232332, E-MTAB-11445, GSE154566, GSE151485, GSE111223, GSE117860, GSE207927, GSE240184, GSE132203, and E-MTAB-17025.

For academic researchers interested in utilizing CheekAge or CheekAI, please reach out to Varun B. Dwaraka. Researchers interested in gaining access to the placebo-controlled clinical trial dataset involving semaglutide should reach out to Michael J. Corley.

## Author Contributions

MNS contributed to computational analysis, project design, writing, and editing. MJC provided the methylomic dataset involving semaglutide. VBD contributed to computational analysis and writing. AAJ took part in project design, writing, and editing.

## Funding

The authors MNS and AAJ are grateful to internal support from Tally Health.

## Conflicts of Interest

MNS and AAJ are former or current employees of Tally Health (New York, New York, United States of America). VBD is a full-time employee of TruDiagnostic (Lexington, Kentucky, United States of America) and MJC serves as a scientific advisor for TruDiagnostic. The authors have no other conflicts of interest or potential conflicts of interest to disclose.

## Abbreviations

AUC: area under the curve
BMI: body mass index
CI: confidence interval
EBP: epigenetic biomarker proxy
FDR: false discovery rate
ICC: intraclass correlation coefficient
MAE: mean absolute error
PC: principal component
PCA: principal component analysis
RMSE: root mean square error
SD: standard deviation
XAI: explainable AI

## References

1. Loyfer N, Magenheim J, Peretz A, Cann G, Bredno J, Klochendler A, et al. A DNA methylation atlas of normal human cell types. Nature. 2023;613(7943):355–64. doi: 10.1038/s41586-022-05580-6.

2. Chen Q, Dwaraka VB, Carreras-Gallo N, Armstrong JF, Sehgal R, Argentieri MA, et al. OMICmAge quantifies biological age by integrating multi-omics with electronic medical records. Nat Aging. 2026;6(3):722–37. doi: 10.1038/s43587-026-01073-7.

3. Gadd DA, Hillary RF, McCartney DL, Zaghlool SB, Stevenson AJ, Cheng Y, et al. Epigenetic scores for the circulating proteome as tools for disease prediction. Elife. 2022;11. doi: 10.7554/eLife.71802.

4. Galkin F, Mamoshina P, Aliper A, de Magalhaes JP, Gladyshev VN, Zhavoronkov A. Biohorology and biomarkers of aging: Current state-of-the-art, challenges and opportunities. Ageing Res Rev. 2020;60:101050. doi: 10.1016/j.arr.2020.101050.

5. Teschendorff AE, Horvath S. Epigenetic ageing clocks: statistical methods and emerging computational challenges. Nat Rev Genet. 2025;26(5):350–68. doi: 10.1038/s41576-024-00807-w.

6. Yu D, Zeng X, Barzilai D, Thor D, Lyu YX. Bridging expectations and science: a roadmap for the future of longevity interventions. Biogerontology. 2025;26(4):138. doi: 10.1007/s10522-025-10278-z.

7. Johnson AA, Shokhirev MN. Understanding and making sense of epigenetic age misalignment across different aging clocks. Epigenomics. 2026;18(8):965–72. doi: 10.1080/17501911.2026.2708567.

8. Johnson AA, Shokhirev MN. Explainable epigenetic aging clocks: an overview of existing AI models and approaches. Expert Rev Mol Diagn. 2026;26(8):695–707. doi: 10.1080/14737159.2026.2693031.

9. Kalyakulina A, Yusipov I, Moskalev A, Franceschi C, Ivanchenko M. eXplainable Artificial Intelligence (XAI) in aging clock models. Ageing Res Rev. 2024;93:102144. doi: 10.1016/j.arr.2023.102144.

10. Kalyakulina A, Yusipov I, Trukhanov A, Franceschi C, Moskalev A, Ivanchenko M. EpInflammAge: Epigenetic-Inflammatory Clock for Disease-Associated Biological Aging Based on Deep Learning. Int J Mol Sci. 2025;26(13). doi: 10.3390/ijms26136284.

11. Shokhirev MN, Johnson AA. Using buccal methylomic data to create explainable aging clocks as well as classifiers and regressors for lifestyle and demographic factors. Front Genet. 2025;16:1637186. doi: 10.3389/fgene.2025.1637186.

12. Prosz A, Pipek O, Borcsok J, Palla G, Szallasi Z, Spisak S, et al. Biologically informed deep learning for explainable epigenetic clocks. Sci Rep. 2024;14(1):1306. doi: 10.1038/s41598-023-50495-5.

13. Milacic M, Beavers D, Conley P, Gong C, Gillespie M, Griss J, et al. The Reactome Pathway Knowledgebase 2024. Nucleic Acids Res. 2024;52(D1):D672–D8. doi: 10.1093/nar/gkad1025.

14. Higgins-Chen AT, Thrush KL, Wang Y, Minteer CJ, Kuo PL, Wang M, et al. A computational solution for bolstering reliability of epigenetic clocks: Implications for clinical trials and longitudinal tracking. Nat Aging. 2022;2(7):644–61. doi: 10.1038/s43587-022-00248-2.

15. Shokhirev MN, Torosin NS, Kramer DJ, Johnson AA, Cuellar TL. CheekAge: a next-generation buccal epigenetic aging clock associated with lifestyle and health. Geroscience. 2024;46(3):3429–43. doi: 10.1007/s11357-024-01094-3.

16. Sehgal R, Markov Y, Qin C, Meer M, Hadley C, Shadyab AH, et al. Systems Age: a single blood methylation test to quantify aging heterogeneity across 11 physiological systems. Nat Aging. 2025;5(9):1880–96. doi: 10.1038/s43587-025-00958-3.

17. Shokhirev MN, Johnson AA. Analysis of variability and epigenetic age prediction across microarray and methylation sequencing technologies. Geroscience. 2025;47(5):6631–8. doi: 10.1007/s11357-025-01824-1.

18. Shokhirev MN, Johnson AA. Various diseases and conditions are strongly associated with the next-generation epigenetic aging clock CheekAge. Geroscience. 2025;47(3):3191–206. doi: 10.1007/s11357-025-01579-9.

19. Clough E, Barrett T. The Gene Expression Omnibus Database. Methods Mol Biol. 2016;1418:93–110. doi: 10.1007/978-1-4939-3578-9_5.

20. Brazma A, Parkinson H, Sarkans U, Shojatalab M, Vilo J, Abeygunawardena N, et al. ArrayExpress--a public repository for microarray gene expression data at the EBI. Nucleic Acids Res. 2003;31(1):68–71. doi: 10.1093/nar/gkg091.

21. Jessen WJ, Borgerding MF, Prasad GL. Global methylation profiles in buccal cells of long-term smokers and moist snuff consumers. Biomarkers. 2018;23(7):625–39. doi: 10.1080/1354750X.2018.1466367.

22. Cruz MAD, Ulfenborg B, Blomstrand P, Faresjo M, Stahl F, Karlsson S. Characterization of methylation patterns associated with lifestyle factors and vitamin D supplementation in a healthy elderly cohort from Southwest Sweden. Sci Rep. 2022;12(1):12670. doi: 10.1038/s41598-022-15924-x.

23. Fitzgerald KN, Hodges R, Hanes D, Stack E, Cheishvili D, Szyf M, et al. Potential reversal of epigenetic age using a diet and lifestyle intervention: a pilot randomized clinical trial. Aging (Albany NY). 2021;13(7):9419–32. doi: 10.18632/aging.202913.

24. Stone TC, Ward V, Hogan A, Ho KMA, Wilson A, McBain H, et al. Using saliva epigenetic data to develop and validate a multivariable predictor of esophageal cancer status. Epigenomics. 2024;16(2):109–25. doi: 10.2217/epi-2023-0248.

25. Katrinli S, Wani AH, Maihofer AX, Ratanatharathorn A, Daskalakis NP, Montalvo-Ortiz J, et al. Epigenome-wide association studies identify novel DNA methylation sites associated with PTSD: a meta-analysis of 23 military and civilian cohorts. Genome Med. 2024;16(1):147. doi: 10.1186/s13073-024-01417-1.

26. Corley MJ, Dwaraka VB, Pang AP, Labbato D, Smith R, Ross Eckard A, et al. Semaglutide slows epigenetic aging in a randomized trial of HIV-associated lipohypertrophy. Nat Commun. 2026;17(1). doi: 10.1038/s41467-026-72861-3.

27. Johnson AA, Sinclair DA. Turning back time: a comprehensive list of interventions that decrease next-generation epigenetic aging clocks in humans. Front Genet. 2026;17:1836446. doi: 10.3389/fgene.2026.1836446.

28. Bischoff-Ferrari HA, Gangler S, Wieczorek M, Belsky DW, Ryan J, Kressig RW, et al. Individual and additive effects of vitamin D, omega-3 and exercise on DNA methylation clocks of biological aging in older adults from the DO-HEALTH trial. Nat Aging. 2025;5(3):376–85. doi: 10.1038/s43587-024-00793-y.

29. Fabian P, Blander G, Deehan R, Torkamani A, Nogal B. Causal impact of genetically-determined fish and fish oil intake on epigenetic age acceleration and related serum markers. Hum Genomics. 2025;19(1):61. doi: 10.1186/s40246-025-00756-3.

30. Bozack AK, Khodasevich D, Nwanaji-Enwerem JC, Gladish N, Shen H, Daredia S, et al. Dietary fatty acids and epigenetic aging in US adults: results from the National Health and Nutrition Examination Survey. NPJ Aging. 2025;12(1):3. doi: 10.1038/s41514-025-00298-x.

31. Lu AT, Quach A, Wilson JG, Reiner AP, Aviv A, Raj K, et al. DNA methylation GrimAge strongly predicts lifespan and healthspan. Aging (Albany NY). 2019;11(2):303–27. doi: 10.18632/aging.101684.

32. Qi W, Gutierrez GE, Gao X, Dixon H, McDonough JA, Marini AM, et al. The omega-3 fatty acid alpha-linolenic acid extends Caenorhabditis elegans lifespan via NHR-49/PPARalpha and oxidation to oxylipins. Aging Cell. 2017;16(5):1125–35. doi: 10.1111/acel.12651.

33. Donega S, Lu AT, Haghani A, Horvath M, Yan Q, Baranzini M, et al. Intermittent hypoxia induces reversible epigenetic age acceleration in old mice. NPJ Aging. 2026;12(1). doi: 10.1038/s41514-026-00425-2.

34. Dwaraka VB, Johnson AA, Seale K, Kahal N, Sheikh D, Morrissey S, et al. Whole-blood DNA-methylation patterns during portable-sauna use in firefighters: an exploratory, single-arm pilot study. bioRxiv. 2026:2026.09.02.748979. doi: 10.64898/2026.09.02.748979.

35. Aryee MJ, Jaffe AE, Corrada-Bravo H, Ladd-Acosta C, Feinberg AP, Hansen KD, et al. Minfi: a flexible and comprehensive Bioconductor package for the analysis of Infinium DNA methylation microarrays. Bioinformatics. 2014;30(10):1363–9. doi: 10.1093/bioinformatics/btu049.

36. Triche TJ, Jr., Weisenberger DJ, Van Den Berg D, Laird PW, Siegmund KD. Low-level processing of Illumina Infinium DNA Methylation BeadArrays. Nucleic Acids Res. 2013;41(7):e90. doi: 10.1093/nar/gkt090.

37. Zheng SC, Breeze CE, Beck S, Dong D, Zhu T, Ma L, et al. EpiDISH web server: Epigenetic Dissection of Intra-Sample-Heterogeneity with online GUI. Bioinformatics. 2019;36(6):1950–1. doi: 10.1093/bioinformatics/btz833.

38. Paszke A, Gross S, Massa F, Lerer A, Bradbury J, Chanan G, et al. PyTorch: an imperative style, high-performance deep learning library. Proceedings of the 33rd International Conference on Neural Information Processing Systems. Curran Associates Inc.; 2019. p. Article 721.

39. Kingma DP, Ba J. Adam: A Method for Stochastic Optimization. 2017 doi: 10.48550/arXiv.1412.6980.

40. Shrout PE, Fleiss JL. Intraclass correlations: uses in assessing rater reliability. Psychol Bull. 1979;86(2):420–8. doi: 10.1037//0033-2909.86.2.420.

41. Bates D, Mächler M, Bolker B, Walker S. Fitting Linear Mixed-Effects Models Using lme4. Journal of Statistical Software. 2015;67(1):1–48. doi: 10.18637/jss.v067.i01.

42. Benjamini Y, Hochberg Y. Controlling the False Discovery Rate: A Practical and Powerful Approach to Multiple Testing. Journal of the Royal Statistical Society: Series B (Methodological). 1995;57(1):289–300. doi: 10.1111/j.2517-6161.1995.tb02031.x.

43. Horvath S. DNA methylation age of human tissues and cell types. Genome Biol. 2013;14(10):R115. doi: 10.1186/gb-2013-14-10-r115.

44. Horvath S, Oshima J, Martin GM, Lu AT, Quach A, Cohen H, et al. Epigenetic clock for skin and blood cells applied to Hutchinson Gilford Progeria Syndrome and ex vivo studies. Aging (Albany NY). 2018;10(7):1758–75. doi: 10.18632/aging.101508.

