## Supplementary Information for "CheekAI: An explainable, deep learning epigenetic aging clock based on lifestyle and health epigenetic biomarker proxies"

#### SUPPLEMENTARY FIGURE LEGENDS

**Supplementary Figure S1. Stability across cross-validation folds.** (a) RMSE, replicate ICC, mean replicate difference, number of significant associations, and longitudinal slope agreement for CheekAI, predicted age (the stage 1 chronological age predictor), and CheekAge in each of the five outer folds (points), with the mean across folds (bars). (b) CheekAI effect per +1 SD of each predicted biomarker in each fold, with that fold's bootstrap 95% confidence interval. A factor that is missing from a fold did not pass that fold's validity filter.

**Supplementary Figure S2. Exploring the effects of semaglutide in a blood dataset.** (a) The impact of semaglutide versus placebo on CheekAI and its components, including various EBPs, in a blood methylomic dataset. (b) CheekAI versus other clocks that were trained using buccal samples in response to semaglutide. These blood samples were derived from semaglutide-treated (n = 45) or placebo-treated (n = 39) patients with HIV-associated lipohypertrophy.

#### SUPPLEMENTARY TABLE LEGENDS

**Supplementary Table S1. Comparison on samples not used to train CheekAge.** The same metrics on all held-out arrays and on the TH arrays, which neither clock was trained on. Longitudinal agreement is platform-corrected, and its p value is from a binomial test against 50%.

**Supplementary Table S2. Sensitivity to the stage-2 selection rule.** CheekAI on held-out people when the stage-2 loss weights are selected under three replicate-reproducibility limits, applied in inner cross-validation. The paper uses the first rule, which was fixed before evaluation.

**Supplementary Table S3. Sample sizes implied by delta-age reliability.** The number of subjects (or tests) needed for 80% power at $\alpha = 0.05$ to detect a true change in delta age. Design 1 is a single-arm pre/post study (subjects), design 2 has two arms with pre/post measurements (subjects per arm), and design 3 is one person tested at evenly spaced times over one year (tests). The ICC and SD are on the delta-age scale (see section 7 of the Extended Methods for additional details).

**Supplementary Table S4. CheekAI in publicly available datasets.** The tests not shown in Figure 4, with the models in section 9 of the Extended Methods. Delta CheekAI and its biomarker part are the group difference (or the change, or the effect per unit of exposure) in years, with its p value in brackets. The strongest biomarker term is the term with the smallest p value, given with its effect in years and, in brackets, its Benjamini–Hochberg FDR across the 21 terms. The value in brackets after the tissue is the correlation of CheekAI with chronological age in the dataset. The note gives the main limitation of each test.

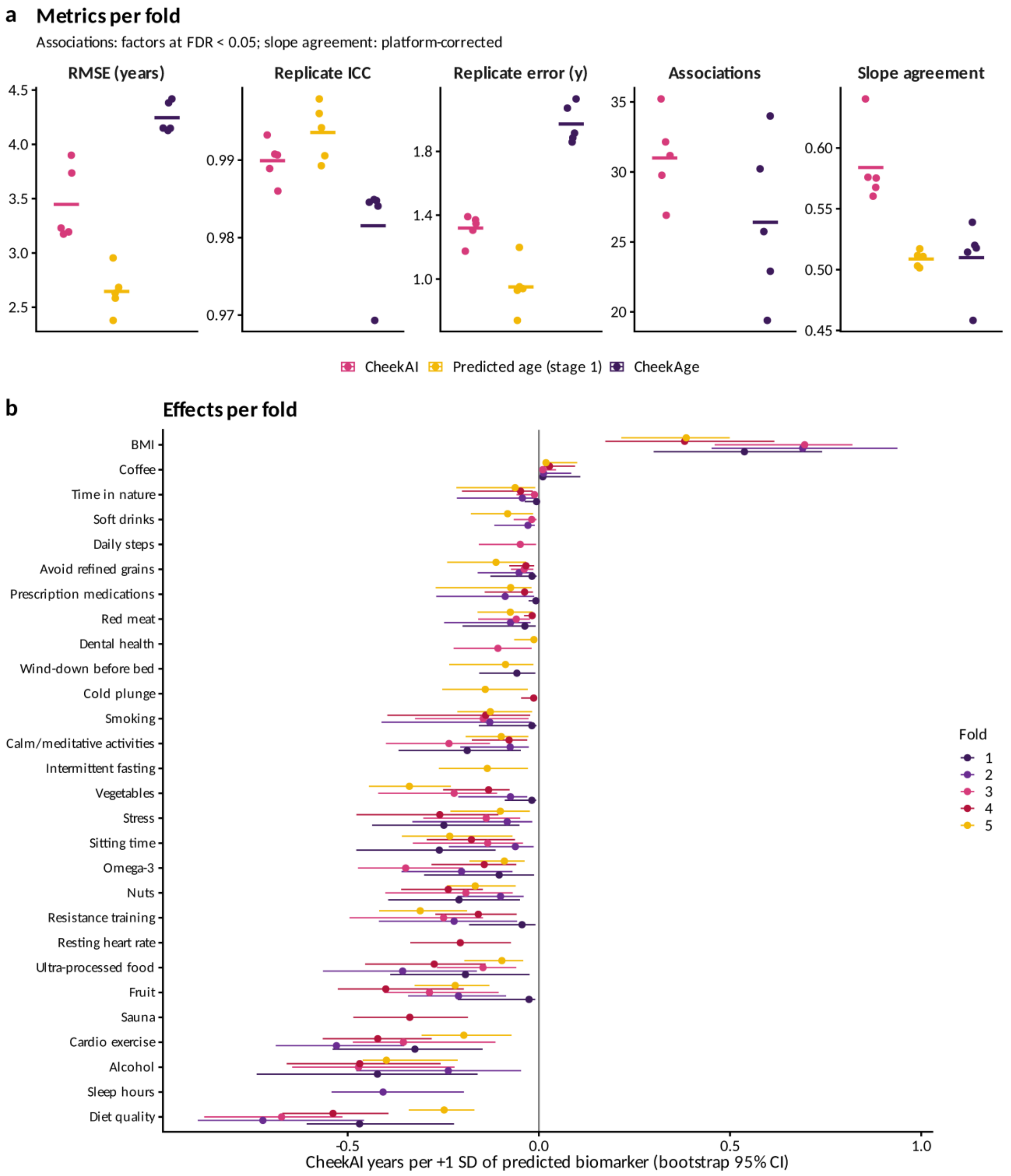
a
Metrics per fold
Associations: factors at FDR < 0.05; slope agreement: platform-corrected
RMSE (years)
Replicate ICC
Replicate error (y)
Associations
Slope agreement
CheekAI
Predicted age (stage 1)
CheekAge
b
Effects per fold
BMI
Coffee
Time in nature
Soft drinks
Daily steps
Avoid refined grains
Prescription medications
Red meat
Dental health
Wind-down before bed
Cold plunge
Smoking
Calm/meditative activities
Intermittent fasting
Vegetables
Stress
Sitting time
Omega-3
Nuts
Resistance training
Resting heart rate
Ultra-processed food
Fruit
Sauna
Cardio exercise
Alcohol
Sleep hours
Diet quality
CheekAI years per +1 SD of predicted biomarker (bootstrap 95% CI)
Fold
1
2
3
4
5

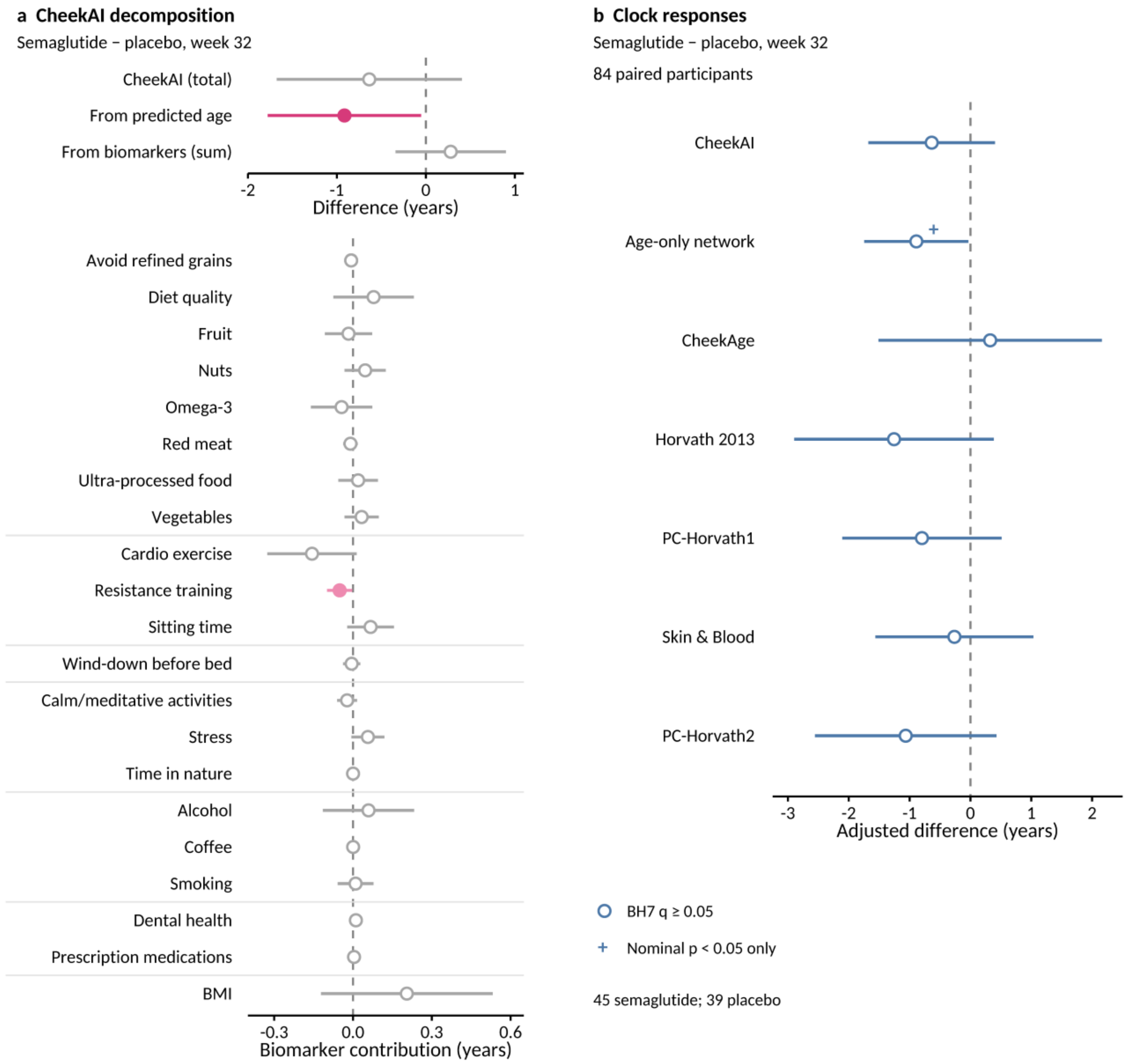
a CheekAI decomposition
Semaglutide − placebo, week 32
CheekAI (total)
From predicted age
From biomarkers (sum)
-2
-1
0
1
Difference (years)
Avoid refined grains
Diet quality
Fruit
Nuts
Omega-3
Red meat
Ultra-processed food
Vegetables
Cardio exercise
Resistance training
Sitting time
Wind-down before bed
Calm/meditative activities
Stress
Time in nature
Alcohol
Coffee
Smoking
Dental health
Prescription medications
BMI
-0.3
0.0
0.3
0.6
Biomarker contribution (years)
b Clock responses
Semaglutide − placebo, week 32
84 paired participants
CheekAI
Age-only network
CheekAge
Horvath 2013
PC-Horvath1
Skin & Blood
PC-Horvath2
-3
-2
-1
0
1
2
Adjusted difference (years)
BH7 q ≥ 0.05
Nominal p < 0.05 only
45 semaglutide; 39 placebo

**Supplementary Table S1.** Comparison on samples not used to train CheekAge.

| Metric | All held-out: CheekAI | All held-out: Predicted age (stage 1) | All held-out: CheekAge | TH only: CheekAI | TH only: Predicted age (stage 1) |
|---|---|---|---|---|---|
| Arrays | 25,400 | 25,400 | 25,400 | 16,915 | 16,915 |
| People | 14,716 | 14,716 | 14,716 | 7,212 | 7,212 |
| r with age | 0.969 | 0.982 | 0.958 | 0.971 | 0.985 |
| MAE (y) | 2.49 | 1.7 | 3.23 | 2.41 | 1.64 |
| RMSE (y) | 3.46 | 2.65 | 4.25 | 3.23 | 2.36 |
| Mean bias (y) | -0.04 | -0.11 | 0.67 | -0.17 | -0.09 |
| Replicate sets | 679 | 679 | 679 | 346 | 346 |
| Replicate ICC | 0.99 | 0.994 | 0.982 | 0.992 | 0.996 |
| Mean replicate \|difference\| (y) | 1.41 | 1.02 | 2.12 | 1.27 | 0.87 |
| Associations at FDR < 0.05 | 39 of 43 | 1 of 43 | 37 of 43 | 34 of 38 | 3 of 38 |
| Mean \|r\| of associations | 0.084 | 0.013 | 0.058 | 0.086 | 0.015 |
| Longitudinal people | 1,676 | 1,676 | 1,676 | 1,317 | 1,317 |
| Longitudinal agreement (%) | 58.5 | 50.8 | 50.9 | 58.7 | 51.3 |
| Longitudinal p | 3.00E-12 | 5.10E-01 | 4.80E-01 | 3.00E-10 | 3.50E-01 |

**Supplementary Table S2.** Sensitivity to the stage-2 selection rule.

| Metric | ICC ≥ 0.99, error ≤ 1.25 y (used) | ICC ≥ 0.9875, error ≤ 1.40 y | ICC ≥ 0.985, error ≤ 1.50 y |
|---|---|---|---|
| RMSE (y) | 3.46 | 3.68 | 4.07 |
| MAE (y) | 2.49 | 2.69 | 3.02 |
| Replicate ICC | 0.99 | 0.989 | 0.986 |
| Mean replicate \|difference\| (y) | 1.41 | 1.5 | 1.69 |
| Associations at FDR < 0.05 | 39 | 39 | 39 |
| Mean \|r\| of associations | 0.084 | 0.09 | 0.098 |
| Longitudinal agreement (%) | 58.5 | 58.7 | 58.9 |
| Longitudinal p | 3.00E-12 | 1.50E-12 | 3.50E-13 |

**Supplementary Table S3.** Sample sizes implied by delta-age reliability.

| clock | icc_delta | sd_delta | sd_error | effect | 1. Pre/post: subjects | 2. Two arms: subjects per arm | 3. One person: tests in 1 year |
|---|---|---|---|---|---|---|---|
| CheekAI | 0.84873988 | 3.37454332 | 1.312433266 | 0.5 | 111 | 218 | 649 |
| CheekAI | 0.84873988 | 3.37454332 | 1.312433266 | 1 | 30 | 56 | 163 |
| CheekAI | 0.84873988 | 3.37454332 | 1.312433266 | 1.5 | 15 | 26 | 73 |
| CheekAI | 0.84873988 | 3.37454332 | 1.312433266 | 2 | 9 | 15 | 41 |
| CheekAI | 0.84873988 | 3.37454332 | 1.312433266 | 3 | 6 | 8 | 19 |
| PredictedAge | 0.85479941 | 2.785279222 | 1.061336077 | 0.5 | 73 | 143 | 425 |
| PredictedAge | 0.85479941 | 2.785279222 | 1.061336077 | 1 | 20 | 37 | 107 |
| PredictedAge | 0.85479941 | 2.785279222 | 1.061336077 | 1.5 | 10 | 17 | 48 |
| PredictedAge | 0.85479941 | 2.785279222 | 1.061336077 | 2 | 7 | 10 | 27 |
| PredictedAge | 0.85479941 | 2.785279222 | 1.061336077 | 3 | 5 | 6 | 13 |
| CheekAge | 0.80904182 | 4.405876839 | 1.925313638 | 0.5 | 235 | 467 | 1397 |
| CheekAge | 0.80904182 | 4.405876839 | 1.925313638 | 1 | 61 | 118 | 350 |
| CheekAge | 0.80904182 | 4.405876839 | 1.925313638 | 1.5 | 28 | 53 | 156 |
| CheekAge | 0.80904182 | 4.405876839 | 1.925313638 | 2 | 17 | 31 | 88 |
| CheekAge | 0.80904182 | 4.405876839 | 1.925313638 | 3 | 9 | 14 | 39 |

**Supplementary Table S4.** CheekAI in publicly available datasets.

| Dataset | Tissue, array (r with age) | Comparison (n) | Delta CheekAI, y (p) | Biomarker part, y (p) | Strongest biomarker term, y (FDR) | Note |
|---|---|---|---|---|---|---|
| GSE94876 | Buccal, 450K (0.79) | Moist snuff users vs non-users (40 vs 40 men) | +0.70 (0.29) | +0.09 (0.76) | Coffee +0.002 (0.77) | CheekAI has no smokeless-tobacco term, and the predicted smokeless-tobacco biomarker did not separate the groups (AUC 0.57); 450K array, 45% of CpGs imputed |
| GSE149747 | Saliva, EPIC (0.77) | Change over 8 weeks within the treated men (19), no control | -0.84 (0.068) | -0.68 (0.017) | BMI -0.294 (0.11) | No control comparison and small; the BMI term fell (p = 0.005) |
| GSE232332 | Saliva, EPIC (0.89) | Non-dysplastic Barrett's oesophagus vs controls (52 vs 115) | +0.26 (0.78) | +1.51 (0.0016) | Fruit +0.186 (0.04) | A precursor condition; the biomarker part was higher (fruit term FDR 0.04) but offset by a lower predicted age |
| E-MTAB-11445 | Saliva, EPIC (0.63) | Alcohol amount per week, per step of a 3-level score (265) | +0.67 (0.24) | +0.62 (0.041) | Nuts +0.070 (0.16) | Coarse questionnaire score; the biomarker part rose with drinking |
| E-MTAB-11445 | Saliva, EPIC (0.63) | Physical activity, per step of a 3-level score (268) | +0.63 (0.17) | +0.51 (0.038) | Smoking +0.079 (0.085) | Opposite to the expected direction; coarse self-report of yearly activity in adults aged 62 to 96 |
| E-MTAB-11445 | Saliva, EPIC (0.63) | Fish and seafood intake, per step of a 3-level score (268) | -0.56 (0.17) | -0.34 (0.12) | Cardio exercise -0.168 (0.11) | Expected direction but not significant; the omega-3 term fell with more fish (p = 0.023, FDR 0.16) |
| GSE154566 | Buccal, EPIC (one age) | Smokers vs non-smokers at age 18 (46 vs 135) | +0.57 (0.24) | +0.05 (0.89) | Coffee +0.007 (3.2e-07) | Little exposure (median 1.3 pack-years among smokers); twins are not independent; 17% of CpGs imputed; predicted smoking AUC 0.58 |
| GSE154566 | Buccal, EPIC (one age) | Per pack-year at age 18 (181) | +0.15 (0.4) | +0.08 (0.54) | Coffee +0.003 (3.7e-07) | As above |
| GSE151485 | Saliva, EPIC (0.97) | Change from before dental surgery to days after short opioid use, visit 1 to 2 (31) | -0.33 (0.46) | -0.59 (0.2) | Ultra-processed food -0.059 (0.48) | Small, no control group, a few days of opioids, and surgery and inflammation change saliva cells |
| GSE151485 | Saliva, EPIC (0.97) | As above, visit 1 to about 6 weeks later, visit 3 (26) | +0.09 (0.86) | +0.25 (0.43) | Nuts +0.089 (0.66) | As above |
| GSE111223 | Saliva, 450K (0.74) | Parkinson's disease vs controls (128 vs 131) | -1.45 (0.031) | -0.16 (0.45) | BMI +0.273 (0.03) | Lower delta CheekAI in patients, from predicted age rather than the biomarkers; 450K array, 45% of CpGs imputed |
| GSE117860 | Blood, EPIC (450K | Current smokers vs non-smokers, men | -0.57 (0.087) | -0.11 (0.36) | Resistance training +0.108 (0.012) | Blood; only the probes shared with the 450K array were deposited |

| | | | | | | |
|---|---|---|---|---|---|---|
| | probes) (0.63) | with HIV (309 vs 220) | | | | (50% of CpGs imputed), and predicted smoking did not separate smokers (AUC 0.51) |
| GSE117860 | Blood, EPIC (450K probes) (0.63) | As above, adjusted for blood cell proportions | +0.06 (0.81) | +0.06 (0.61) | Dental health +0.007 (0.2) | As above |
| GSE207927 | Blood, EPIC (0.64) | Per kg/m$^2$ of BMI, 2017 wave (108) | +0.02 (0.83) | +0.10 (0.024) | Diet quality +0.030 (0.29) | Blood; the biomarker part rose with BMI. The 2007 wave was not analysed because CheekAI did not track age in it (r = -0.05) |
| GSE240184 | Blood, EPIC (0.95) | Change over 3 months of behavioural weight loss (62), no control | -0.04 (0.82) | -0.13 (0.33) | Prescription medications +0.003 (0.17) | Blood; no control group, and weight change was not deposited |
| GSE132203 | Blood, EPIC (0.93) | Current PTSD vs none (745) | +0.89 (0.0067) | +0.08 (0.58) | Smoking +0.033 (0.78) | Blood; the difference came from predicted age, not the biomarkers |
| GSE132203 | Blood, EPIC (0.93) | As above, adjusted for blood cell proportions | +0.71 (0.018) | +0.05 (0.72) | Smoking +0.036 (0.73) | As above |
| GSE132203 | Blood, EPIC (0.93) | Moderate or severe childhood abuse vs none (786) | +0.53 (0.058) | -0.05 (0.71) | Diet quality -0.109 (0.13) | Blood; as for PTSD, from predicted age |
| GSE132203 | Blood, EPIC (0.93) | Per type of lifetime trauma (702) | +0.05 (0.27) | -0.01 (0.64) | Fruit -0.007 (0.85) | Blood |
| E-MTAB-17025 | Blood, EPIC (0.96) | Change over 1 month, vegan vs meat-rich diet (48) | +0.68 (0.094) | +0.30 (0.28) | Smoking +0.105 (0.82) | Blood, small and short; the deposited diet labels disagree with the assigned diet for 20 of 48 people |

### EXTENDED METHODS

#### 1. Study population and sample metadata

The initial analysis set comprised 25,403 arrays. Three lacked chronological age and were excluded, leaving 25,400 arrays from 14,716 people.

*Replicates, people and timepoints*

Arrays marked as replicates in the metadata (the same swab or person assayed more than once at one collection) share a replicate identifier. There were 679 replicate sets with 1,750 arrays; 390 sets had more than two arrays and 556 spanned more than one platform. Repeat collections from the same person are linked in the metadata by a pointer to the previous sample. People were defined as the connected components of the graph that joins replicate arrays and previous-sample links (union–find), which gave 14,716 people. Timepoints are the distinct collection ages of a person after replicate arrays are collapsed. In total, 3,402 people had at least two timepoints, 2,220 had at least three, and 2,036 had at least three timepoints spanning more than six months (maximum 12; median follow-up 0.92 years among people with at least two timepoints).

*Biomarkers*

Fifty variables besides chronological age were modeled. The survey items covered diet (diet quality, ultra-processed food, fruit, red meat, refined grains, vegetables, nuts, artificial sweeteners, omega-3, intermittent fasting, and poultry), physical activity (resistance training, cardio exercise, sitting time, daily steps, and overall activity), sleep (sleep quality, sleep hours, a wind-down routine before bed, and caffeine timing), mind and social (stress, social connection, sense of purpose, engagement with calm/meditative activities, and time spent in nature), substances (cigarette smoking, alcohol, soft drinks, smokeless tobacco, coffee, tea, and e-cigarettes), recovery (sauna and cold plunge), health (dental health, a recent injury/operation, prescription medications, hazardous work exposure, resting heart rate, sickness, self-rated health, and self-perceived aging), body size (BMI), demographics (sex, race/ethnicity, education, and menopause age), and technical (log median unmethylated signal, log median methylated signal, and estimated epithelial fraction). The epithelial cell fraction (Epi) was estimated with EpiDISH [1] while the log median methylated and unmethylated signal intensities (mMed and uMed) came from minfi [2].

Survey questionnaires changed over time, so items were answered by between 1,935 and 18,635 samples. Answers were coded so that higher values (closer to 1) are healthier. BMI values below 8 were set to missing. Values between 8 and 16 were taken to be weights entered in kilograms where pounds were expected and were multiplied by 2.20462, which was confirmed against the recorded height and weight. Sex was coded 1 for female.

*Overlap with the CheekAge training set*

CheekAge was trained on 8,045 samples from an earlier subset of this cohort [3]. Most of them are among the 8,485 arrays whose barcode does not start with "TH". The remaining 16,915 arrays, from 7,212 people, have TH barcodes and were never used to train CheekAge. Barcodes were attached to the training rows by matching each array's median methylated and unmethylated signals, which are unique to each array.

**2. DNA methylation data**

Raw IDAT files were processed with minfi (v1.54.1) using noob background and dye-bias correction [4]. For EPIC v2, probe suffixes were stripped and the first probe was kept for duplicated CpGs. The analysis matrix contained the 244,860 CpGs measured on all three platforms (EPIC v1, EPIC v2, and a custom EPIC array), stored as single-precision beta values. At inference, missing beta values in external data are imputed with the training-set mean of each CpG.

**3. Principal component representation**

The beta matrix $X$ (samples × CpGs) was represented by an exact uncentered principal component decomposition, as in the original model. It was computed from the sample Gram matrix:

$$G = XX^T$$

This Gram matrix was accumulated in double precision over blocks of CpGs. For any set of training samples, the eigendecomposition of the corresponding block of $G$ gives the left singular vectors $U$ and the singular values $d$, and the leading 10,000 components were kept. Training samples are represented by their rows of $U$. A new sample $x$ is projected out of sample as

$$u = x\,V\,\text{diag}(d)^{-1}, \quad V = X^T U\,\text{diag}(d)^{-1}$$

or, equivalently, $u = G_{\text{new, train}}\,U\,\text{diag}(d)^{-2}$. This makes it possible to refit the PCA inside every cross-validation training set without materializing the full matrix, so that held-out samples never inform the components.

Eigendecompositions used double-precision LAPACK (scipy.linalg.eigh) or the equivalent GPU routine (PyTorch linalg.eigh), depending on the available memory. Because the trailing components are nearly degenerate, the two solvers can return slightly different bases, so model training is not bit-for-bit deterministic across hardware. The released model therefore fixes its PCA by shipping the loadings $V$ and the singular values $d$ with SHA-256 checksums. Projecting the training samples through the released loadings reproduces their predictions to a relative error of $5.5 \times 10^{-8}$.

**4. Stage 1: Biomarker networks**

*Architecture*

Each of the 51 biomarkers has its own network, implemented in PyTorch [5], that takes the 10,000 PCs as input. The survey and demographic biomarkers use a standard network with the layers Linear(10,000 → 128), ReLU, Linear(128 → 64) and Linear(64 → 1). The five continuous, high-precision targets (chronological age, BMI, Epi, mMed and uMed) use a large network with the layers Linear(10,000 → 512), ReLU, Linear(512 → 256), ReLU, Linear(256 → 256) and Linear(256 → 1). All biomarkers within a class share the same architecture.

*Loss*

For biomarker $b$, the loss is a weighted mean squared error over the samples where $b$ is observed, plus a replicate-agreement penalty:

$$L_b = \frac{1}{N_b} \sum_i \omega_i \, (\hat{y}_i - y_i)^2 + 0.2 \cdot \overline{|\hat{y}_i - \hat{y}_j|}$$

The bar denotes the mean over the pairs $(i, j)$ of technical-replicate arrays in the batch. Discrete biomarkers (fewer than 20 levels) are weighted by inverse class frequency, $\omega_i = N/(C \cdot N_c)$, where $C$ is the number of classes and $N_c$ the number of samples in the class of sample $i$; for all other biomarkers $\omega_i = 1$. Mini-batches of 2,048 samples are built so that all arrays of a replicate set fall in the same batch, and gradients are clipped to a norm of $1000/\sqrt{\text{epoch} + 1}$.

*Optimization and hyperparameter search*

Networks were trained with Adam [6], with the learning rate decayed exponentially per epoch. Each biomarker had its own grid search on inner fold 0 (see Section 6 of the Extended Methods), over the settings below.

| Class | Decay γ | Max epochs | Patienc e | Learning rates | Weight decay | Configurations |
|---|---|---|---|---|---|---|
| Standard networks | 0.992 | 1,000 | 100 | 1e-4, 3e-4, 1e-3, 3e-3, 1e-2 | 1e-4, 1e-3, 1e-2, 5e-2 | 21 (incl. original 5e-3/1e-3) |
| Large networks | 0.998 | 5,000 | 300 | 3e-4, 1e-3, 3e-3, 1e-2 | 1e-4, 1e-3, 5e-3, 2e-2, 5e-2 | 21 (incl. original 1e-2/5e-3) |

Standard networks start their output bias at the training mean. The setting and the stopping epoch were chosen by validation correlation for the survey biomarkers, because for these weak ordinal traits the loss barely distinguishes a predictive model from one that predicts the mean, and by validation loss for the large networks. Refits use the chosen setting for the chosen number of epochs.

*Collapsed fits*

A fit whose predictions on its own training samples vary by less than 5% of the trait's observed SD has collapsed to a near-constant output, typically under strong weight decay; for comparison, a biomarker that passes the validity filter (r ≥ 0.1) has a calibrated spread of at least 10% of the SD. Collapsed fits were treated as training errors. Early stopping only selects epochs at which the network has not collapsed, and collapsed settings are ineligible in tuning. If any inner or outer refit with the chosen setting collapses, that setting is excluded for that biomarker in that fold, the next-best eligible setting is selected, and all of that biomarker's refits are redone, so that each biomarker uses one setting per fold. In the released model this happened for four biomarkers (sickness, fruit, calm or meditative activities, and smokeless tobacco), and none of the final networks are collapsed.

*Re-centering and numerical precision*

After training, each network's output bias is set so that its mean prediction on its own training samples equals their mean observed value. This removes per-refit offsets that depend on the stopping point; it uses training data only and does not change the network's ranking of samples. Some strongly regularized survey networks predict within a very narrow range (a spread as small as about $10^{-7}$ around a value near the trait mean), which is below single-precision resolution once the output bias is added. The final linear layer, the bias and all later calibration are therefore evaluated in double precision.

An adaptive SGD optimizer used in earlier versions of this model was compared with Adam in a pilot on fold-1 training people. It gave no consistent gain, so Adam was used throughout.

**5. Stage 2: CheekAI adjustment**

*Expected directions*

For each lifestyle or health factor $f$, the expected direction $s_f \in \{-1,0,+1\}$ is the sign of its Pearson correlation with CheekAge acceleration in the training people if the Benjamini–Hochberg FDR is below 0.05, and $s_f = 0$ otherwise. CheekAge values are those recorded in the production database, CheekAge acceleration is the residual of CheekAge on chronological age, and the direction is oriented so that the unhealthy direction raises epigenetic age. There were 43 candidate factors. Chronological age was excluded, as were race/ethnicity and education (which by design never adjust age), the technical and cellular variables, sex and menopause age. Medication count keeps its empirical direction (more medications, older).

*Validity filter and scale*

Stage 2 uses out-of-fold stage-1 predictions for the training people, produced by refitting every network on three inner folds and predicting the fourth. Each refit's predictions are expressed in units of that refit's own training-set distribution, as (prediction − training mean) / SD of its predictions on its training samples. This is needed because networks that move little from their initial output differ between refits far more in center and spread than in signal (spreads up to about 200-fold apart), so pooling raw predictions, or transferring a rescaling estimated on the inner refits to the outer refit, would be dominated by refit artefacts. For well-behaved networks this standardization changes almost nothing, and networks whose predictions do not vary beyond double-precision rounding contribute zero. A biomarker is valid if its pooled out-of-fold prediction correlates with the observed value at $r \geq 0.1$ with FDR < 0.05, and only valid lifestyle and health biomarkers enter stage 2. Valid biomarkers are rescaled by variance matching, so that the pooled out-of-fold predictions have the observed mean and SD of the trait. Composed with the outer refit's own scale, this gives one fixed linear calibration per biomarker, which is stored in the model.

*Model*

With predicted age $\hat{A}$ and standardized predicted biomarkers $z_f$ (standardized with the training means and SDs),

$$\text{CheekAI} = \hat{A} + \sum_{f:\, s_f \neq 0} s_f \ \text{softplus}(\theta_f)\, z_f$$

where $\text{softplus}(\theta_f) = \ln(1 + e^{\theta_f}) \geq 0$ is the effect of factor $f$ in years per SD and $\theta_f$ is initialized at −3.

*Loss*

For a training set with chronological age $A$ and CheekAI age $C$, let the delta age be $\Delta = C - A$ and the age acceleration $\Delta_{\text{adj}}$ be $\Delta$ residualized on $A$ by least squares. The loss is

$$L = w_{\text{mse}}\, \overline{\Delta^2} + w_{\text{rep}}\, R^2 + w_{\text{assoc}}\, (1 - S)^2 + w_{\text{long}}\, (1 - \rho)^2$$

Here $R$ is the mean absolute difference of $C$ between consecutive arrays of each replicate set, and $S = \frac{1}{F} \sum_f s_f \ \text{corr}(y_f, \Delta_{\text{adj}})$ is the mean signed correlation of the observed factors $y_f$ with age acceleration, each computed over the samples where $f$ is observed. $\rho$ is the correlation across people between two per-person slopes over time, the slope of $\Delta_{\text{adj}}$ and the slope of the lifestyle index. Per-person slopes are ordinary least-squares slopes against age at collection, for people with at least three timepoints with a lifestyle index spanning more than 0.5 years. The lifestyle index is the mean of a sample's observed factors after z-scoring and orienting each by $s_f$, so that higher values are less healthy.

*Weight grid and training*

The loss weights were searched over 60 settings: $w_{\text{mse}} \in \{0.01, 0.001\}$, $w_{\text{rep}} \in \{1, 4\}$, $w_{\text{assoc}} \in \{4, 16, 32, 64, 128\}$ and $w_{\text{long}} \in \{2, 4, 8\}$. Each setting was trained full-batch with Adam (learning rate 0.003, weight decay 0.001) for up to 4,000 epochs, with early stopping (patience 300) on the inner validation loss.

*Pre-specified selection rule*

For each outer fold, every setting was evaluated by 4-fold inner cross-validation over people. Settings with a mean inner-validation one-way replicate ICC of at least 0.99 and a mean replicate absolute error of at most 1.25 years were eligible. Among the eligible settings, the one maximizing the mean signed association plus the mean slope correlation ($S + \rho$, computed on held-out inner people) was chosen, and it was refit on all outer-training people for its mean early-stopping epoch count. The longitudinal weight is never zero; its range is small because a stronger longitudinal push trades off against replicate reproducibility. For the released model the chosen setting was $w_{\text{mse}} = 0.001$, $w_{\text{rep}} = 4$, $w_{\text{assoc}} = 64$ and $w_{\text{long}} = 4$ (1,265 epochs), with 21 valid factors. Supplementary Table S2 shows the held-out results under two looser limits.

An alternative monotone network adjustment (non-negative weights with monotone activations) was compared with the additive form on inner-validation samples of folds 1 and 2. It produced implausibly lopsided extreme adjustments without better validation metrics, so the additive form was used for all folds.

*Effect estimates*

The effect of factor $f$ is reported as the change in CheekAI per +1 training SD of the predicted biomarker, $s_f$ softplus$(\theta_f)$. The 95% confidence intervals are percentile intervals over 100 bootstrap refits of stage 2 that resample people with replacement and hold the stage-1 predictions fixed. Because the effects are sign-constrained, the intervals cannot cross zero, so their width is the informative quantity.

**6. Cross-validation design**

People (not arrays) were randomly assigned to 5 outer folds (seed 123), so all replicate arrays and timepoints of a person are in the same fold. Within each outer-training set, people were split into 4 inner folds (seed 124 + k for outer fold k). For each outer fold, the stage-1 settings were first tuned by fitting the PCA on inner folds 1–3 and evaluating the grid on inner fold 0. Out-of-fold predictions were then produced by refitting the PCA and all 51 networks with the chosen settings on three inner folds and predicting the fourth, in turn, which gave out-of-fold biomarker predictions for every training person. The PCA and all 51 networks were next refitted on all outer-training people. Validity, rescaling and the stage-2 weight selection were done by inner cross-validation on the out-of-fold predictions, and the chosen stage-2 model was refit on all training people. Finally, the held-out fold was projected onto the outer PCA and predicted by the assembled model.

Held-out people are used only for evaluation. Pooled results concatenate the five held-out folds, so each person is evaluated exactly once by a model that never saw them. The released model repeats the same steps, except the evaluation, with all 25,400 samples as the training set (inner folds seeded 129).

#### 7. Evaluation metrics

All metrics are computed on held-out people, pooled over the five outer folds unless stated otherwise.

*Accuracy*

Accuracy is summarized by the correlation, MAE, RMSE and mean bias of CheekAI and of predicted age against chronological age.

*Replicate reproducibility*

The one-way random-effects intraclass correlation [7] is computed from the first two arrays of each replicate set as

$$\text{ICC}(1) = \frac{MS_B - MS_W}{MS_B + MS_W}$$

together with the absolute differences between those arrays (mean, median and the percentage below 0.5 years), split by same and different platform. Reliability on the delta-age scale was estimated with a linear mixed model in lme4 [8]: predicted age is modeled as a natural cubic spline of chronological age plus a random intercept per replicate set. The spline degrees of freedom (1–6) are chosen by AIC under maximum likelihood, the model is refit by REML, and ICC $= \sigma_b^2/(\sigma_b^2 + \sigma_e^2)$. The same model gives the sample-size calculations in Supplementary Table S3.

*Age-adjusted associations*

Age acceleration is the residual of (clock − chronological age) on chronological age, fitted within each held-out fold so that calibration differences between folds stay out of the residuals. For each of the 43 candidate factors, the Pearson correlation of age acceleration with the observed factor is computed, with Benjamini–Hochberg FDR across factors. Significant associations are counted overall and in and against the expected direction. The expected direction comes from training people; it never differed in sign between folds, so each factor has one consensus direction.

*Longitudinal agreement*

This applies to the 1,676 people with at least three timepoints with a lifestyle index spanning more than 0.5 years. For each person, two ordinary least-squares slopes over age at collection are computed, the slope of delta age and the slope of the observed lifestyle index. Agreement is the fraction of people whose two slopes share a sign, tested with a two-sided binomial test against 50%. Because arrays from different platforms can carry systematic offsets, the platform-corrected version subtracts the mean delta age of each platform, estimated in training people, before the slopes are computed.

*Biomarker prediction*

For each biomarker, the held-out Pearson correlation between predicted and observed values is averaged over folds.

**8. Comparison with CheekAge**

CheekAge was computed on the same arrays with the platform-matched published model: Tx100Mini for EPIC v1, Tx100r2mini for EPIC v2 and Tx250kmini for the custom array. These per-platform values reproduce the CheekAge values recorded in the production database ($r = 0.995$ over the 23,245 arrays with a database value; median absolute difference 0.55 years). All CheekAge metrics are computed exactly as for CheekAI, on the same held-out samples.

Most of CheekAge's training samples are among the 8,485 held-out arrays without a TH barcode (see Section 1 of the Extended Methods), which favors CheekAge in the main comparison. Every metric was therefore repeated on the 16,915 TH arrays (Supplementary Table S1). Age acceleration keeps its fold-specific fit. Replicate metrics use the 346 replicate sets whose arrays are all TH, and longitudinal agreement uses the 1,317 people all of whose arrays are TH. Five factors were answered by fewer than 30 TH samples and were not tested, which left 38 factors for the associations.

**9. Publicly available, external datasets**

*Selection*

GEO [9] and ArrayExpress [10] were searched for DNA methylation datasets from buccal cells, saliva, or blood that were measured on the Illumina 450K or EPIC array and had at least 50 samples and a lifestyle exposure, an intervention, or a disease recorded for each sample. Twelve datasets were analyzed: GSE94876, E-MTAB-11445, GSE149747, GSE232332, GSE117860, GSE207927, GSE240184, GSE132203, GSE154566, GSE151485, GSE111223, and E-MTAB-17025.

*Data processing*

For GSE111223, GSE132203, GSE151485, GSE232332, GSE240184, E-MTAB-11445, and E-MTAB-17025, the IDAT files were processed with minfi noob as for the training data (see Section 2 of the Extended Methods), in batches of 100 arrays. For the other datasets the deposited beta values were used: GenomeStudio average beta values for GSE94876 (450K) and GSE149747, ssNoob-normalized beta values for GSE117860, of which only the probes shared by the 450K and EPIC arrays were deposited, the authors' normalized beta values for GSE154566, and the processed beta values deposited for GSE207927. CpGs missing from a dataset were set to their training means, as the package does for any input. The median fraction of CheekAI's CpGs filled in this way was 45% for the two 450K datasets (GSE94876 and GSE111223), 50% for GSE117860, 17% for GSE154566 and at most 2.3% for the others. CheekAI was predicted for all datasets.

*Outcomes*

Delta CheekAI is the residual of CheekAI on chronological age within the dataset; in GSE154566, where every sample was taken at age 18, it is CheekAI minus its mean. The predicted age part is the same residual for predicted chronological age, and the biomarker part is CheekAI minus predicted chronological age, which is the sum of the 21 biomarker terms $s_f w_f z_f$ (see Section 5 of the Extended Methods). Each of these outcomes, and each of the 21 terms, was regressed on the exposure with the covariates given below, and the coefficient of the exposure is reported in years. Because every model is linear and includes age where age varies, the coefficient for delta CheekAI equals the sum of the coefficients for its two parts, and the coefficient for the biomarker part equals the sum of the coefficients for the 21 terms. Benjamini–Hochberg FDR was computed across the 21 terms within each test. Single predicted biomarkers were also checked directly against the matching observed variable, by the area under the ROC curve for two groups or by Pearson correlation for a continuous variable.

*Models*

In GSE94876, cigarette smokers and moist snuff users were each compared with non-tobacco users, adjusted for age. In E-MTAB-11445, the questionnaire answers on alcohol amount per week, yearly physical activity, as well as fish and seafood intake were coded as scores (low 0, intermediate 1, high 2), drinking frequency per week was coded as high (1) against low (0) with the intermediate answers left out, and every model was adjusted for age and sex; smoking was not tested because only four participants smoked. In GSE149747, each person's change from before to after the 8-week programme was regressed on treatment, and the change within the treated individuals alone was tested by the intercept. In GSE232332, the replicate arrays of each person were averaged, and cancer (invasive oesophageal adenocarcinoma, intramucosal cancer and high-grade dysplasia) and non-dysplastic Barrett's oesophagus were each compared with controls (healthy volunteers and people with no positive diagnosis at endoscopy), adjusted for age and sex. In GSE117860, current smokers were compared with non-smokers, adjusted for age, and in a sensitivity analysis also for the deposited proportions of CD8 T cells, CD4 T cells, natural killer cells, B cells and monocytes. In GSE207927, BMI (per kg/$m^2$) was tested in the 2017 samples, adjusted for age and sex. The 2007 samples were not analyzed because CheekAI did not track chronological age in them ($r = -0.05$, against 0.64 in 2017), so the change between the two waves could not be used either. In GSE240184, the change from baseline to 3 months was tested by the intercept. In GSE132203, current post-traumatic stress disorder, moderate or severe childhood abuse and the number of types of lifetime trauma were each tested with adjustment for age and sex, and post-traumatic stress disorder also with the deposited blood cell proportions. In GSE154566 (buccal swabs at age 18), smokers (any pack-years) were compared with non-smokers, and pack-years were tested as a continuous exposure, both adjusted for sex; the twins were analyzed as independent samples. In GSE151485, the change from visit 1 (before dental surgery) to visit 2 and to visit 3 was tested by the intercept. In GSE111223, Parkinson's disease was compared with controls, adjusted for age, sex and ethnicity. In E-MTAB-17025, the change over one month on a vegan diet was compared with the change on a meat-rich diet, adjusted for the deposited binary diet column, which disagrees with the assigned diet for 20 of the 48 people and most likely marks the two cohorts of the study.

**10. Software and computation**

Models were trained with Python 3.11, PyTorch 2.13 (CUDA 13.0), NumPy 2.4, SciPy 1.17 and pandas 3.0 on a single NVIDIA Quadro RTX 5000 GPU (16 GB). Statistical analyses and figures used R 4.5.0 with lme4 2.0.6, minfi 1.54.1, ggplot2 and cowplot, and the figures were exported as editable vector graphics with officer and rvg. The prediction package takes beta values (CSV, TSV or Parquet, or IDAT files through the included minfi script) and returns CheekAI, predicted chronological age, the 51 predicted biomarkers and the per-biomarker contributions in years. The same prediction is available as a step-by-step R script that needs no deep-learning library; on the example sample it agrees with the Python package to within $10^{-5}$ years.

**References**

1. Zheng SC, Breeze CE, Beck S, Dong D, Zhu T, Ma L, et al. EpiDISH web server: Epigenetic Dissection of Intra-Sample-Heterogeneity with online GUI. Bioinformatics. 2019;36(6):1950-1. doi: 10.1093/bioinformatics/btz833.
2. Aryee MJ, Jaffe AE, Corrada-Bravo H, Ladd-Acosta C, Feinberg AP, Hansen KD, et al. Minfi: a flexible and comprehensive Bioconductor package for the analysis of Infinium DNA methylation microarrays. Bioinformatics. 2014;30(10):1363-9. doi: 10.1093/bioinformatics/btu049.
3. Shokhirev MN, Torosin NS, Kramer DJ, Johnson AA, Cuellar TL. CheekAge: a next-generation buccal epigenetic aging clock associated with lifestyle and health. Geroscience. 2024;46(3):3429-43. doi: 10.1007/s11357-024-01094-3.
4. Triche TJ, Jr., Weisenberger DJ, Van Den Berg D, Laird PW, Siegmund KD. Low-level processing of Illumina Infinium DNA Methylation BeadArrays. Nucleic Acids Res. 2013;41(7):e90. doi: 10.1093/nar/gkt090.
5. Paszke A, Gross S, Massa F, Lerer A, Bradbury J, Chanan G, et al. PyTorch: an imperative style, high-performance deep learning library. Proceedings of the 33rd International Conference on Neural Information Processing Systems. Curran Associates Inc.; 2019. p. Article 721.

6. Kingma DP, Ba J. Adam: A Method for Stochastic Optimization. 2017 doi: 10.48550/arXiv.1412.6980.
7. Shrout PE, Fleiss JL. Intraclass correlations: uses in assessing rater reliability. Psychol Bull. 1979;86(2):420-8. doi: 10.1037//0033-2909.86.2.420.
8. Bates D, Mächler M, Bolker B, Walker S. Fitting Linear Mixed-Effects Models Using lme4. Journal of Statistical Software. 2015;67(1):1 - 48. doi: 10.18637/jss.v067.i01.
9. Clough E, Barrett T. The Gene Expression Omnibus Database. Methods Mol Biol. 2016;1418:93-110. doi: 10.1007/978-1-4939-3578-9_5.
10. Brazma A, Parkinson H, Sarkans U, Shojatalab M, Vilo J, Abeygunawardena N, et al. ArrayExpress--a public repository for microarray gene expression data at the EBI. Nucleic Acids Res. 2003;31(1):68-71. doi: 10.1093/nar/gkg091.
